# Partial Correlation as a Tool for Mapping Functional-Structural Correspondence in Human Brain Connectivity

**DOI:** 10.1101/2024.10.16.618230

**Authors:** Francesca Santucci, Antonio Jimenez-Marin, Andrea Gabrielli, Paolo Bonifazi, Miguel Ibáñez-Berganza, Tommaso Gili, Jesus M. Cortes

## Abstract

Brain structure-function coupling has been studied in health and disease by many different researchers in recent years. Most of the studies have estimated functional connectivity matrices as correlation coefficients between different brain areas, despite well-known disadvantages compared to partial correlation connectivity matrices. Indeed, partial correlation represents a more sensible model for structural connectivity since, under a Gaussian approximation, it accounts only for direct dependencies between brain areas. Motivated by this and following previous results by different authors, we investigate structure-function coupling using partial correlation matrices of functional magnetic resonance imaging (fMRI) brain activity time series under various regularization (a.k.a. noise-cleaning) algorithms. We find that, across different algorithms and conditions, partial correlation provides a higher match with structural connectivity retrieved from Density Weighted Imaging data than standard correlation, and this occurs at both subject and population levels. Importantly, we also show that regularization and thresholding are crucial for this match to emerge. Finally, we assess neuro-genetic associations in relation to structure-function coupling, which presents promising opportunities to further advance research in the field of network neuroscience, particularly concerning brain disorders.

## 1 Introduction

A fundamental problem in network neuroscience is understanding the relationship between **Functional Connectivity** (FC), accounting for similarity in the activation patterns between brain areas, and **Structural Connectivity** (SC), which maps the brain’s anatomical connections ([1, 2, 3, 4, 5, 6, 7, 8, 9, 10, 11, 12, 13, 14, 15, 16, 17, 18]). Numerous, rapidly evolving functional states emerge from the relatively static structural connectivity ([7]). The underlying structure partially determines function and activity that, in turn, shapes the structure through the processes of neuromodulation and plasticity ([7, 19, 20]). The investigation of the relationship between structure and function as a biomarker is generally referred to as structure-function coupling (SFC) ([21]).

It has been found that, while the intensities (*weights*) of the single connections (*links*) of the structural and functional connectivity, are positively correlated at rest ([5]), this correlation is not always consistent and exhibits variability across individuals ([22, 23]), age ([19, 23]), cognitive tasks ([22]), brain regions ([24, 23]), and in brain-related disorders ([25, 26, 27, 28, 29]).

Nevertheless, the relationship between SC and FC does not follow a simple mapping at the level of the single link. For instance, two regions can be functionally connected without a direct structural connection, and SC evolves over much longer time scales than FC ([20]). For this reason, the SC-FC correspondence has been investigated at the module or aggregate level, exploring the full set of nested partitions within a hierarchical tree, revealing that the structural and functional connectivities share a common modular architecture ([30, 31, 9, 32, 33]).

FC is usually estimated as the correlation matrix between pairs of time series of activity (usually blood oxygen level-dependent (BOLD) functional magnetic resonance imaging (fMRI) signals at rest) from different brain areas, and, less commonly, as the partial correlation (PC) ([34, 35, 36, 37]).

Several advantages of PC over correlation have been shown. For instance, PC represents a more direct model for brain connectivity, since, under a linear (Gaussian) approximation, it accounts for direct dependencies between brain areas only ([38, 39, 40, 41, 42]) and, accordingly, it provides a higher link-wise match between SC and FC ([42], see also S4 in Supplementary Information (SI) for a more detailed description of the relation between Structure and Function in the linear approximation). Moreover, PC-based FC exhibits reduced variance across subjects ([43]) and yields higher prediction scores for certain individual-level measures ([36, 44]).

However, the use of the PC is limited by the low accuracy of its statistical estimation in the small-sample limit, that is when the time series are short relative to the number of brain regions considered, which is often the case with BOLD fMRI time series ([40, 42, 41, 45]). To address this issue, various regularization methods have been proposed within network neuroscience for accurate inference of the correlation and precision matrices ([35, 40, 43, 36]), many of which introduce *ℓ*_1_ or *ℓ*_2_ penalty terms. The *ℓ*_1_ penalty leads to sparse estimators, such as the Graphical Lasso (GLASSO) ([46]), while the penalty *ℓ*_2_ results in Linear Shrinkage (LS) estimators, where matrices are not intended to be sparsified ([47, 48, 41]). The GLASSO is, in general, a good approach for structure recovery ([49]), but it may not yield stable covariance coefficients. Moreover, its costly optimization is not suited for large-scale datasets. On the other hand, the Linear Shrinkage estimator is simple and fast to compute. It yields biased estimates that are more stable than the empirical covariance and is often recommended for functional connectivity ([36]).

These methods have been compared in terms of non-imaging features prediction power ([36, 44, 37]), stability across scans ([50, 36, 37]), retrieval of synthetic data generative models ([51, 40, 37]), and measures such as test-set (or out-of-sample) likelihood ([52, 45]), suggesting that the choice between sparse or shrinkage estimators may depend on the intended use and interpretation of FC.

Indeed, the dependence of the SFC on the FC inference method, including PC, has already been investigated ([37, 42]), but only at the single links level, while it has been suggested that the SFC may be rooted in a common hierarchical modular organization rather than in a correspondence between the single connections ([30, 9, 33, 31]).

Here, we go beyond both approaches by exploring the correspondence between SC and FC at the hierarchical aggregated level, inferring FC from both correlation and partial correlation matrices. Specifically, we show that: i) FC, when estimated from the regularized PC, exhibits greater similarity to SC in terms of modular structure, both at the subject and at the population level, and a smaller variance across subjects; ii) regularization is crucial to our results, which are robust with respect to the regularization methods most widely used in neuroscience; iii) sparsity is fundamental for the emergence of the FC’s hierarchical modular structure; iv) the SC-FC similarity reaches a maximum in correspondence with a specific partition of FC; v) the partition’s modules are characterized by neurogenetic expression present in major diseases.

These findings advance our understanding of SFC in the brain, indicating that the use of *regularized PC matrices* may provide a more accurate and stable representation of SC, enhancing the potential for clinical applications and insights into neurogenetic disease mechanisms.

## 2 Results

### 2.1 Connectivity matrices and cross-modularity for assessing the SC-FC correspondence

**SC graph** 𝒢_*S*_ and **FC graph** 𝒢_*F*_ (or connectomes) were obtained from data previously published in ([31]) and available at ([53]) for a population of *P* = 136 healthy participants and *N* = 183 regions of interest (ROIs) (see Fig. 1 for a scheme of the methodological pipeline). In particular, we used diffusion-weighted imaging (DWI) matrices to extract a unique estimate of SC for each subject. To estimate FC, we used resting state (rs) fMRI signal of *T* = 652 time steps (Fig. 1a). In particular, we use two differentiated strategies to estimate FC: from the correlation **C** and from the partial correlation 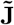 matrices, respectively. The latter is defined as 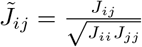, or the standardized version of the **precision matrix J** = **C**^*−*1^. In the fMRI time series, cast as a *N* × *T* matrix **X**, the ratio *q* = *N/T* is not negligible, so *regularization* methods are needed to accurately estimate **C** (and even more its inverse **J** ([45])) beyond the **sample correlation matrix** 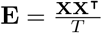(the time series will be assumed from now on to be demeaned and standardized). Here, we estimated such **regularized correlation matrix C**_*µ*_ and **regularized partial correlation matrix** 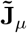 according to different regularization algorithms *µ* (Fig. 1b and Methods).

**Figure 1.**
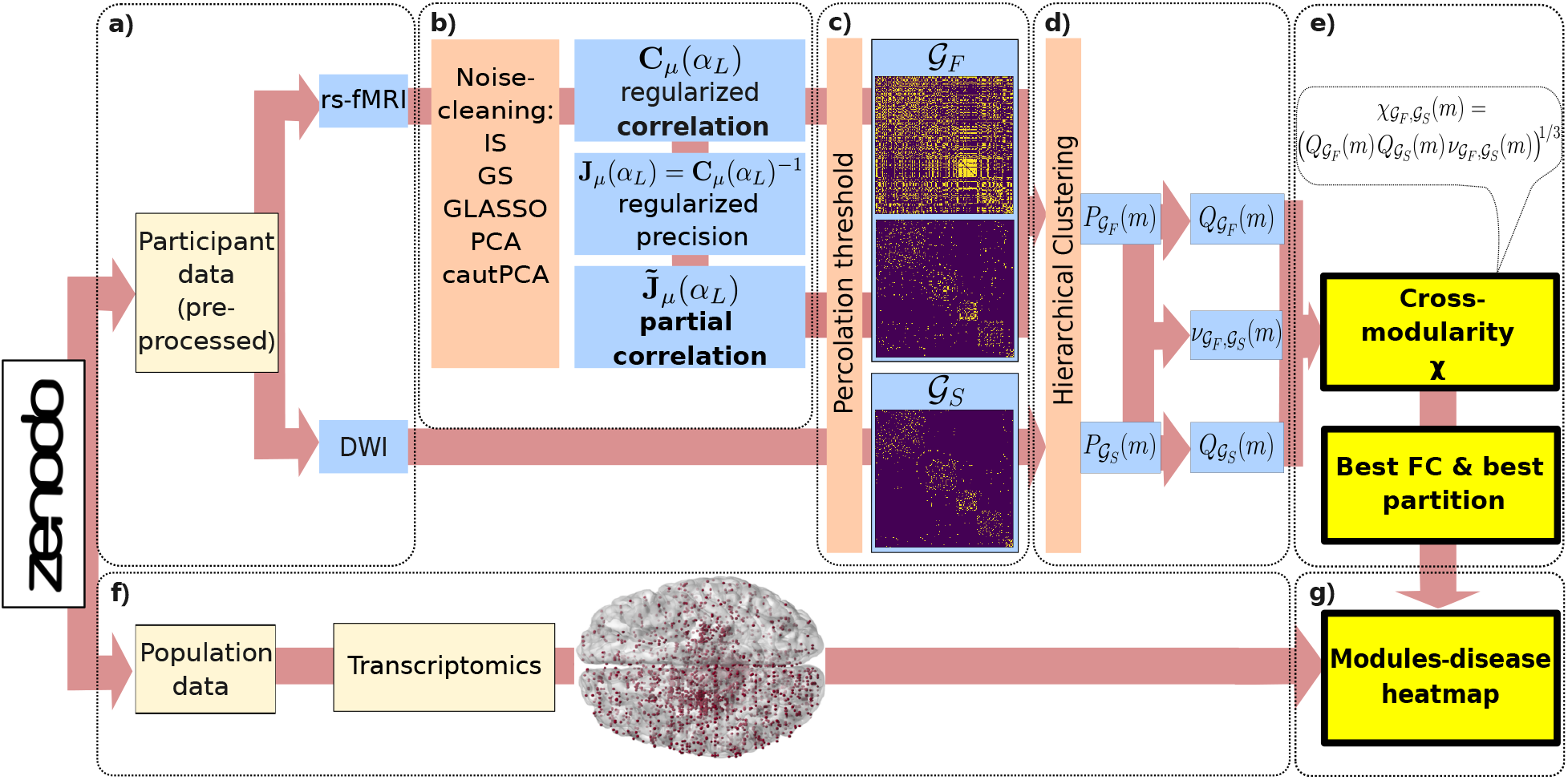
Methodological sketch and pipeline. **a)** Preprocessed data of 136 healthy subjects have been obtained from the open dataset ([31]), at ([53]) Data include, for each subject, a Diffusion Weighted Imaging (DWI) matrix of *N* = 183 brain regions, and rs-fMRI BOLD time series of *N* brain regions and *T* = 652 time steps. **b)** Each subject FC is estimated as i) the regularized correlation matrix **C**_*µ*_(*α*_*L*_) of the time series, where *µ* stands for the specific regularization method, and *α*_*L*_ for the parameter that maximizes the validation-set likelihood, given the method, or ii) as the regularized partial correlation matrix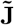, the standardized version of regularized precision matrix (the correlation inverse) 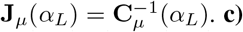 We cut both functional and structural matrices, taken in absolute value, at the percolation threshold. These thresholded matrices are the adjacency matrices of the sparse graphs 𝒢_*F*_ and 𝒢_*S*_. **d)** We computed the hierarchical clustering of 𝒢_*F*_ and 𝒢_*S*_, separately. This computation returns two sets of nested partitions 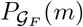 and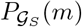, in *m* modules, for *m* = 2, …, *N*. Subsequently, for each couple of partitions 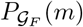 and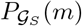, we computed their individual quality 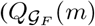 and 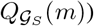and their agreement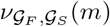. **e)** These three measures allow us to compute the cross-modularity 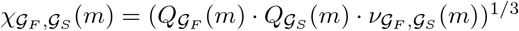, whose maximum can be used to identify a meaningful partition of the population FC. **f)** We made use of transcriptomic data extracted from ([31]) and data relative to the association between genes and a set of diseases extracted from ([54]) to (**g**) evaluate the association between each of the modules of the retrieved FC partition and these diseases.

Next, we construct three connectivity matrices **M** for each subject, one for the SC (containing the DWI data) and two for FC: *M*_*ij*_ = |*C*_*ij*_| and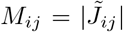, with | · | indicating the absolute value. Afterwards, we cut the connectivity matrices at the so-called *percolation threshold* ([55]). This procedure prescribes setting to zero all matrix elements smaller than the cutoff that would break the corresponding graph in two or more connected components, so that the thresholded graphs’ density depends on their topological properties (also see methods). Therefore, interpreting these thresholded matrices as adjacency matrices, SC and FC graphs were generated (Fig. 1c). In the following, we will indicate the structural connectivity graph as 𝒢_*S*_ and the functional connectivity graph as 𝒢_*F*_, possibly distinguishing between the graphs inferred from **C** or 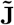 as 𝒢_*F*_ (**C**) and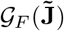, respectively. If not otherwise specified, we assume all graphs to be at the percolation threshold. Each pair of SC and FC graphs was then compared at the module level. This was done for each subject in the dataset, and at the population level as well. To this aim, we performed the hierarchical clustering of both 𝒢_*S*_ and 𝒢_*F*_, separately. This procedure returns two independent sets of nested partitions into *m* modules, 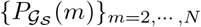 and 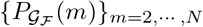. For all values of *m* = 2, …, *N* we computed: i) the quality 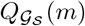of 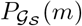 and the quality 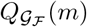 of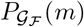, in terms of Newman’s modularity ([56]), and ii) the agreement, 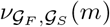, between the pair of partitions 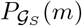 and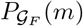, in terms of adjusted Normalized Mutual Information (NMI) ([57]). Notice that, here, we did not maximize *Q* but simply calculated it at all levels in the hierarchical clustering (Fig. 1d). These three quantities allowed us to compute the **cross-modularity**

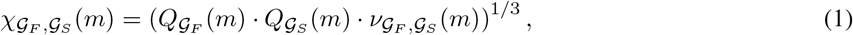

which is a slightly different measure from the original one defined in ([30]). The aim of cross-modularity is to quantify the reciprocal similarity of a couple of graphs in terms of their hierarchical modular structure while also taking into account the quality of their individual partitions. This is done for all number of modules, so that a complete comparison of the whole hierarchy of nested partitions of the two graphs is provided. The range of *χ*(*m*) follows straightforwardly from those of Newman modularity and NMI: they both reach a maximum of 1 for a high-quality partition and a perfect match and are expected to vanish in case of random and unrelated partitions ([58, 59, 57, 60, 61]). In this work, cross-modularity is always used to compare 𝒢_*S*_ with an estimate of 𝒢_*F*_ therefore, when referring, for the sake of shortness, to the “cross-modularity of 𝒢_*F*_,” we will always mean the cross-modularity of between 𝒢_*F*_ and 𝒢_*S*_ in Eq. (1).

The cross-modularity *χ*(*m*) was then used for i) comparing the effects of estimating 𝒢_*F*_ through **C** or through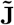, as well as the impact of regularization, in terms of similarity between the hierarchical nested structures of 𝒢_*F*_ and 𝒢_*S*_ at both the individual and population levels; ii) extracting a meaningful partition of our estimate of population-level FC graph (Fig. 1e). Finally, we made use of transcriptomic data (Fig. 1f) ([31]) to better motivate the biological interpretation of the different estimated partitions (Fig. 1g).

### 2.2 Partial correlation enhances the FC-SC correspondence in the human brain

As represented in Fig. 2, 𝒢_*F*_ is more similar to 𝒢_*S*_ when inferred from regularized partial correlation than when inferred from regularized correlation, in terms of hierarchical modularity. This result is robust across all regularization strategies that are most common in neuroscience. More in detail, with reference to Fig. 2a, we found that the cross-modularity *χ*(*m*) of 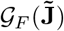 was much lower than the one of 𝒢_*F*_ (**C**) when no regularization was applied. However, when regularization was applied, 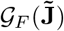 reached higher values of *χ*(*m*) for most regularization methods. On the other hand, the cross-modularity of 𝒢_*F*_ (**C**) did not exhibit significantly different variations after regularization (as expected). Moreover, 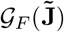 exhibited a significantly higher *χ*(*m*) than 𝒢_*F*_ (**C**), for methods such as Identity shrinkage (IS), Group Shrinkage (GS), Graphical Lasso (GLASSO), and Cautious PCA (cautPCA). In addition, FC inferred from partial correlation exhibited a smaller across-subject variance. A similar trend was found measuring the spectral distance (Fig. 2b) between each couple of subjects’ matrices (see Methods). We additionally observed that the cross-modularity of 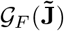 is higher when regularizing with the method cautPCA ([45]), than when regularizing with PCA (alternatively known as *Eigenvalue Clipping* ([62])), of which cautPCA is a variant.

**Figure 2.**
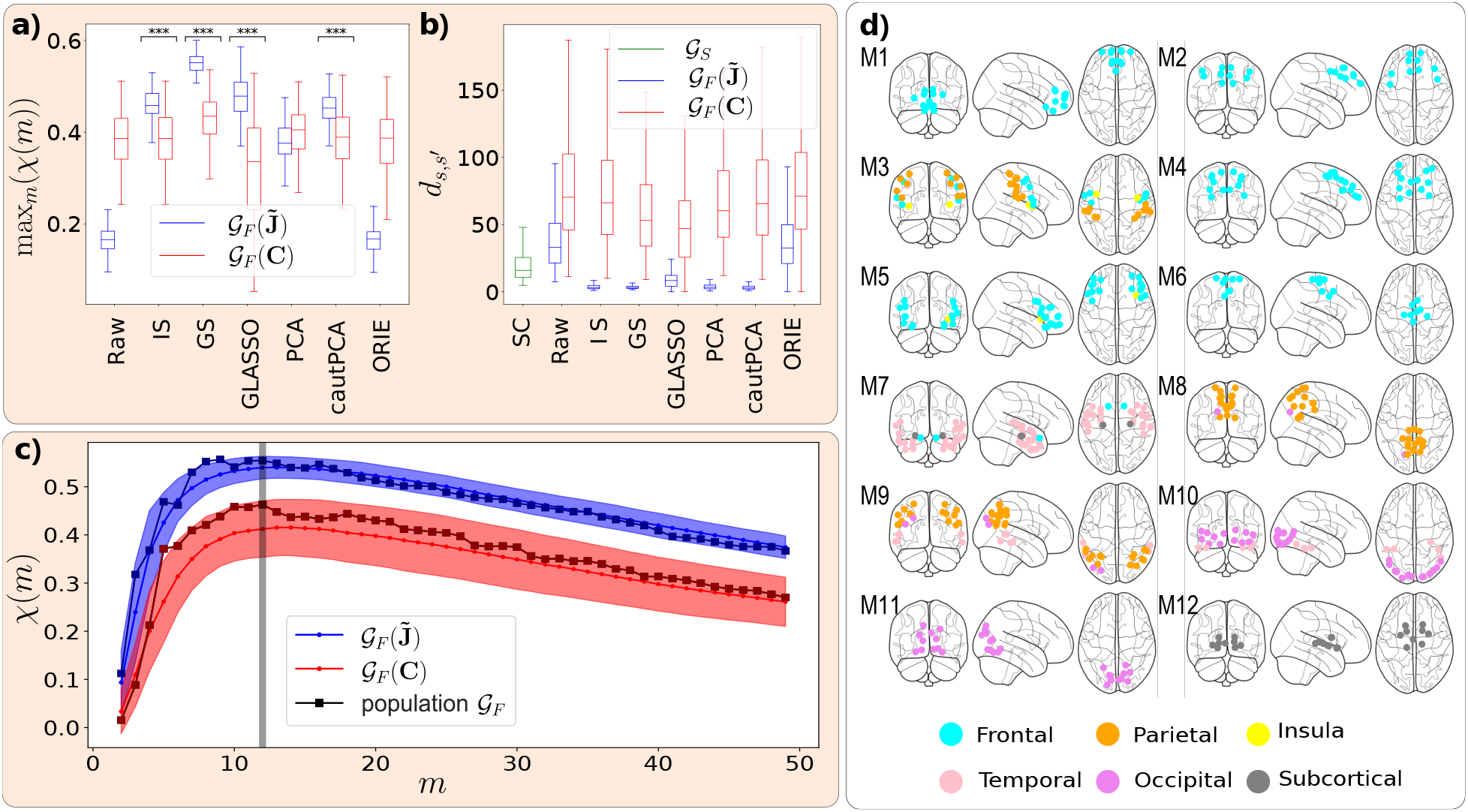
Partial correlation enhances the FC-SC correspondence in the human brain. Panels **a** and **b** show how two relevant measures such as (**a**) the cross-modularity and (**b**) the across-subject variance, measured in terms of spectral distance, change depending on whether 𝒢_*F*_ is estimated as **C** (red) or 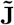(blue), and the relevance of using a regularization method (“raw” stands for no regularization). More in detail, the boxplots represent the across-subject distribution of (**a**) max_*m*_ *χ*(*m*) and (**b**) the spectral distances between each couple of subjects’ 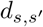. Most regularization strategies (denoted with three asterisks) significantly enhance the cross-modularity of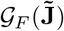. Moreover, the subjects’ 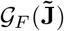 exhibit a significant and systematic lower spectral distance, with respect to 𝒢_*F*_ (**C**), similarly to 𝒢_*S*_, that we have reported in green for comparison. We have assessed such significance by performing a Mann-Whitney U rank test of the null hypothesis that the distributions underlying the pair of samples (relative to **C** and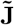) are the same. More in detail, in **a** the methods marked by the asterisks are associated with a p-value smaller than 10^*−*3^, and the alternative hypothesis is that theto **C** is stochastically *less* than the distribution underlying the sample corresponding to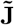. On the other hand, in **b** all methods are associated with a vanishing p-value, and the alternative hypothesis is that the distribution underlying the sample corresponding to **C** is stochastically *greater* than the distribution underlying the sample corresponding to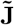. The cross-modularity curves of 𝒢 _*F*_ (**C**) and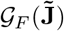, regularized with GS, are shown in subfigure **c**, both at the subject (the solid lines are the across-subject means and the shaded areas their standard-deviation) and at the population level (square-solid lines). The curves of 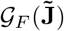are significantly higher than those of 𝒢_*F*_ (**C**) for almost all values of *m* and are characterized by a lower variance. The grey vertical line indicates the number of modules *m*^*\**^ = 12 that maximizes the cross-modularity of most subjects. We reported in subfigure **d** the partition of the population 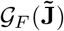into *m*^*\**^ modules, drawing the ROIs belonging to each module as dots in the brain glass plots, colored according to the anatomical regions.

### 2.3 Cross-modularity curves identify representative partitions

The complete cross-modularity curves for 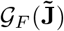 and 𝒢_*F*_ (**C**), both for individual subjects and at the population level are shown in Fig. 2c. These curves are obtained using the GS regularization method, as the one providing the highest PC cross-modularity, but we found qualitatively consistent results across regularization methods. We found that, in general, at both the individual subjects and population levels, for _*F*_ inferred from both regularized **C** and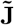, and across all regularization methods, the cross-modularity *χ*(*m*) increases rapidly with the number of modules *m* when *m* is low. It exhibits a soft maximum when *m* is between 10 and 20 (though its exact position may slightly vary depending on the specific method used for inferring FC) and then decreases slowly as *m* increases. We stress that, since the cross-modularity is the product of two quantities that are expected to vanish in unstructured ([59, 58]) or unrelated graphs ([61, 60]), the fact that it presents a maximum is nontrivial, and the number of modules providing it can be used as a criterion for selecting the number of modules to partition the FC. Accordingly, Fig. 2d presents the hierarchical clustering partition of 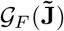 into 12 modules (the median across-subject argument of max_*m*_ *χ*(*m*)) of the population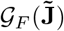.

See the SI for further details regarding the FC-SC correspondence at the population level (Fig. S3), and the overlap between each module and the Desikan-Killiany structural areas and the Yeo functional networks (Figs. S4c-d).

### 2.4 Robustness of the enhanced FC-SC similarity with respect to the regularization parameters

In this article, we understand *regularization* in the sense of statistical inference: the regularized estimator leads to a higher out-of-sample data likelihood, at the expense of a lower training-set likelihood ([45]). The regularized estimator presents, in other words, a lower variance error with respect to the maximum likelihood estimator, at the expense of a higher bias error (and a lower bias+variance error). We now present an assessment of the robustness of our results with respect to the value of the regularization parameter, generically referred to as *α* for all the regularization methods. The regularization parameter is such that, for all the methods except ORIE, *α* = 0 represents no regularization (the maximum-likelihood estimator), while *α* = 1 represents the completely biased estimator. Given a subject time series **X**, we fix the value of *α*_L_ as the one that maximizes the validation-set likelihood **X**_val_, given the method (see Sec. Methods).

We have already seen (see Fig. 2a) that regularization is crucial for our estimation of PC-based FC to exhibit an enhanced similarity to SC. We now show that, actually, the enhancement of FC-SC cross-modularity induced by statistical regularization is very close to *its maximum possible value*, understood as the maximum value of the FC-SC cross-modularity over all values of *α*. Let us define *α*_*χ*_ as the value of the regularization parameter that maximizes the (PC-based) FC-SC cross-modularity (given a regularization method). We observe (see Fig. 3), that the effect of *statistical regularization* (i.e., with parameter *α*_L_) in terms of FC-SC similarity is, rather remarkably, very similar to the effect of regularization methods with parameter *α*_*χ*_, despite the statistical regularization does not use any information about the SC matrix. The histograms for both parameters *α*_*χ*_ and *α*_L_ are, however, clearly different (see Fig. 3). We conclude that regularization is crucial to observe the enhancement of FC-SC similarity, and that such enhancement is, at the same time, robust with respect to the regularization parameter *α*, as far as *α* is of the same order of *α*_L_ (not orders of magnitude lower).

Remarkably, these results remain qualitatively identical in a backup analysis with a supplementary extra dataset (see Fig. S2 in the SI).

**Figure 3.**
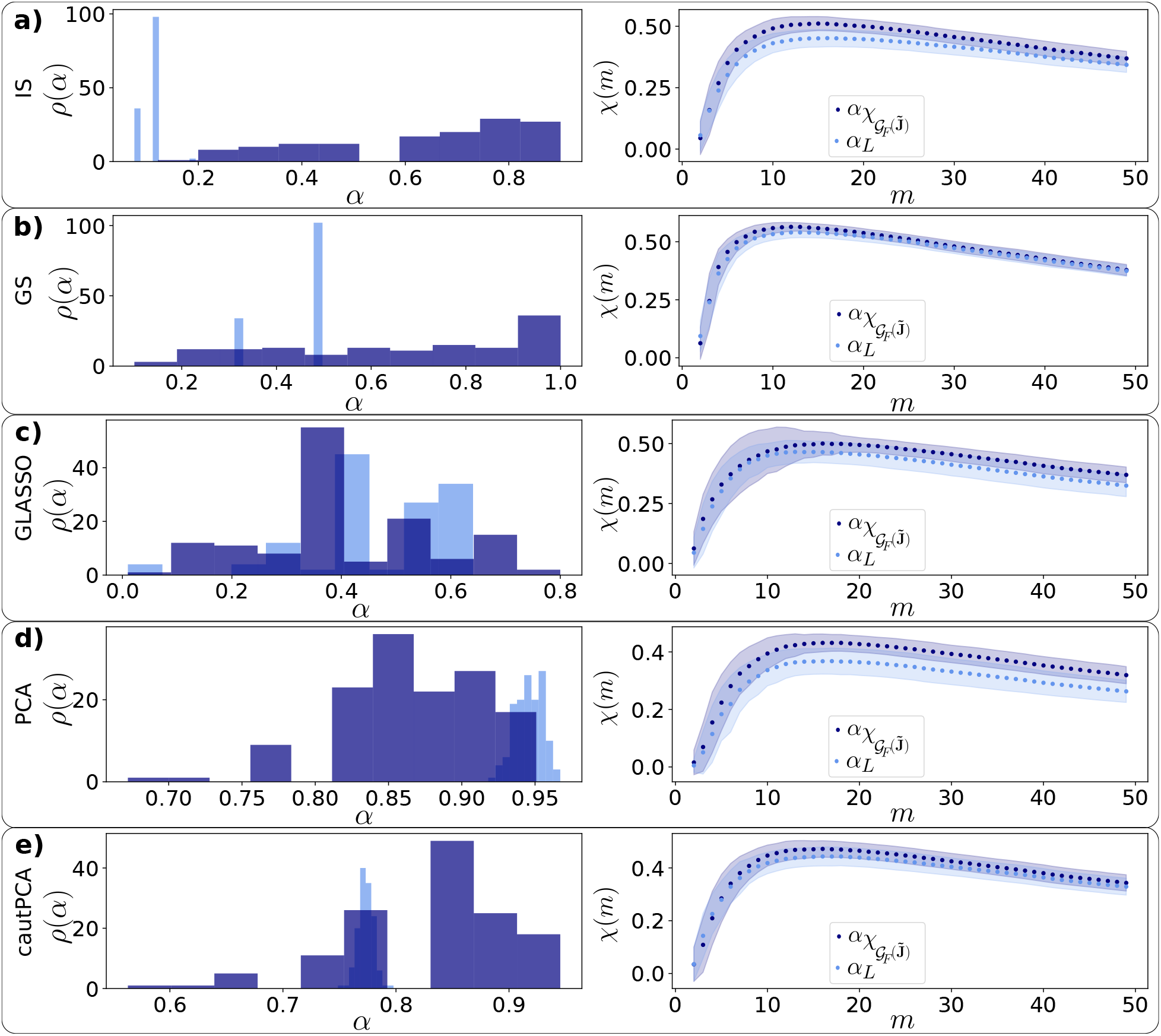
Robustness of the enhanced FC-SC similarity with respect to the regularization parameters, for different regularization methods (rows **a-e**). First column: across-subjects distribution of the regularization parameter *α*_*L*_ that maximizes the validation-set loglikelihood (light blue) and the parameter 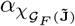 that maximizes the peak of the cross-modularity (*χ* = max_*m*_ *χ*(*m*)) of 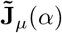 (dark blue), given the regularization method *µ*. Although the distributions of *α*_*L*_ and 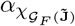 are different, they produce similar *χ* curves: the second column represents the cross-modularity curves of 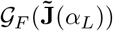(light blue) and of 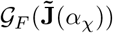 (dark blue), where the dotted lines are the across-subject means and the shaded areas represent the plus/minus one-standard-deviation distance from the mean.

### 2.5 Thresholding is crucial for the partial correlation FC-SC correspondence

We observed that a high similarity between 𝒢_*S*_ and 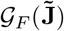only emerged provided that the latter was inferred from sparse partial correlation matrices (see Fig. 4a). It is noteworthy that thresholding disrupts the positive definiteness of the matrices, rendering the thresholded connectivity matrices non-compliant with the mathematical definition of correlation matrices. Consequently, some researchers, including [41], advocate against the thresholding process. Instead, they recommend the sparsification induced by GLASSO to achieve sparsity, when needed. Indeed, we have confirmed that 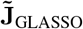is a sparse matrix and that further thresholding 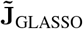 up to the percolation threshold, only slightly enhances the cross-modularity of 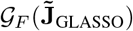(see Fig. 4b). The same does not apply to **C**_GLASSO_, which is a dense matrix (Fig. 4c).

**Figure 4.**
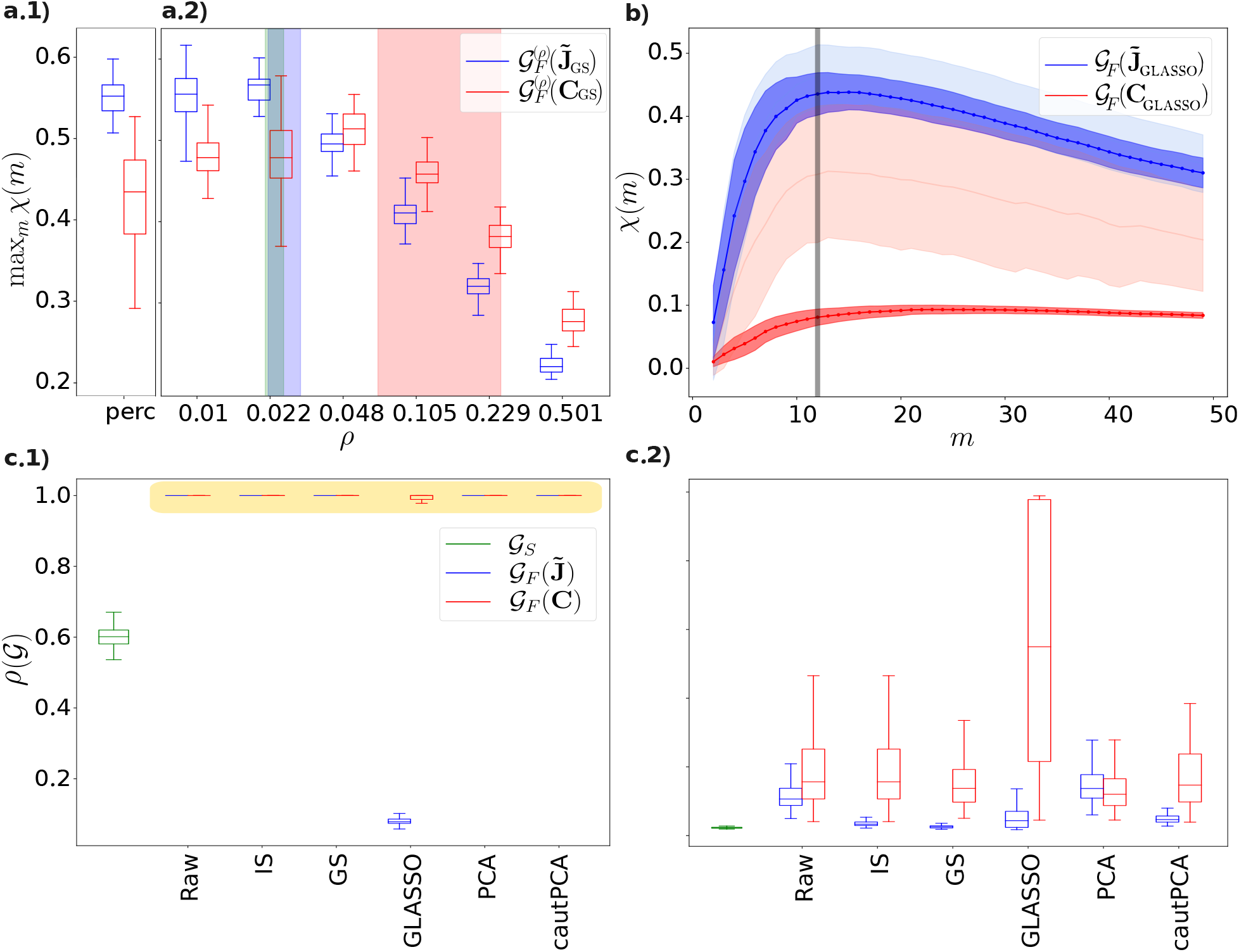
Relevance of thresholding. Subfigures **a.1**,**a.2** report the dependency of cross-modularity on the FC density *ρ*: the boxplots in **a.2** represent the maximum (across modules) FC-SC cross-modularity, for the partial correlation-based FC 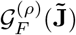(blue) and the correlation-based FC 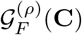 (**C**) (red), regularized with GS and cut at density *ρ* (while the SC graph is always cut at percolation). As a comparison, we show in **a.1** (“perc”) the values that are obtained if all graphs are cut at the percolation threshold (the same as in Fig. 2). We also report the average across subjects, plus/minus one standard deviation, of the percolation density of: structural connectivity, PC-based FC, correlation-based FC (green, blue, and red vertical thick bars, respectively). These trends are robust across regularization methods. Anyway, notice that GLASSO returns a PC matrix that is much sparser than the corresponding correlation matrix (as shown in subfigure **c.1**). This is why in this case, if no threshold is applied, PC provides significantly higher cross-modularity curves with respect to the correlation: subfigure **b** shows the cross-modularity curves for **C** (red) and 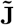(blue), regularized with GLASSO, without any threshold (darker curves) and with percolation threshold (lighter curves) as a comparison. The boxplots **c.1, c.2** show the across-subject distribution of the single subjects’ densities *ρ*(𝒢) of the SC (green) and FC graphs obtained from regularized **C** (red) and 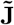 (blue) matrices, without (**c.1**) and with (**c.2**) the cut at the percolation threshold, for different regularization strategies (*“Raw”* stands for “no regularization”). We observe how only the partial correlation regularized with GLASSO provides an FC graph that is already sparse (note that the symbols highlighted in yellow in subfigure **c.1** appearing as horizontal lines are, in fact, zero-error boxplots).

### 2.6 Relevance of FC graphs’ density

It is well known that a graph’s density is very relevant for many graph properties ([63]). Therefore, in this section we address the question of whether the higher FC-SC match in terms of partial correlations is attributable to the mere fact that the partial-correlation-based FC is sparser.

To answer this question, we estimated the FC-SC match in terms of cross-modularity, but *as for fixed values of the FC network density ρ* (tuned through thresholding), *equal for correlation- and PC-based FC* (while all SC graphs are cut at percolation). If 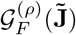 exhibits a significantly higher match than 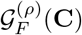 in a wide range of common values of *ρ*, we can conclude that the density is not the determinant factor of the phenomenon that we describe.

The results of this analysis are shown in Fig. 4a.2, reporting the maximum (across modules) of the FC-SC modularity for different densities (in abscissa) of the FC graphs constructed from the regularized **C** and 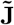matrices. For each subject in Fig. 4a.2, we regularize the **C**, 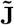matrices with the GS method; we then construct the associated graphs’ adjacency matrices, and threshold them up to the x-axis density; we finally compute the across-subjects values of *χ*, reported in the box-plots. The average percolation density, plus and minus one standard deviation, is represented in the same figure, as the three vertical bars colored in green, blue and red for the SC, the partial correlation-based FC and correlation-based FC, respectively. For ease of comparison, we also include in Fig. 4a.1 the cross-modularity for the case in which all the matrices are cut at the so-called (subject-dependent) percolation threshold (labeled “perc”). On the one hand, we observe that sparser graphs, whose density *ρ* is more similar is to the SC density *ρ*_*χ*_, tend to present higher FC-SC match. On the other hand, we observe that, interestingly, the 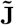-based match is significantly higher than the **C**-based match, for a wide range of sufficiently low values of *ρ*. This implies that we cannot simply attribute the higher PC-based FC-SC match to the fact that PC-based FC is sparser (hence, more similar in density to the SC density).

The results of Fig. 4**-a.2** are qualitatively identical when we threshold the SC at the same density as the FC and when we use different regularization criteria.

### 2.7 Partition of the population FC and neurogenetic interpretation

Finally, we characterized the modules of the population 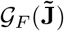partition shown in Fig. 2 in terms of their participation in major brain-related disorders. In other words, we assessed whether some modules exhibit a significantly lower or higher expression of genes associated with a particular disease. To this end, we computed the across-modules Z-score of the median transcriptomic expression values for sets of genes associated with each brain-related disorder across the ROIs in each module, following a procedure similar to that described in ([31]). The results, shown in Fig. 5, reveal that module M12, corresponding to subcortical regions, is highly relevant for most disorders. It is positively associated with tumor-related and neurodegenerative diseases and negatively associated with psychiatric, substance abuse, and movement-related ones. We additionally checked the stability of the population 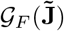 partition across 100 bootstrap samplings of the dataset. Indeed, we found an overall average NMI similarity score of 0.86 out of 1 between the main partition and those computed at each sampling (see methods). We also found that some modules are particularly persistent: notably, module M12 appears identical in all partitions (Fig. 5b).

**Figure 5.**
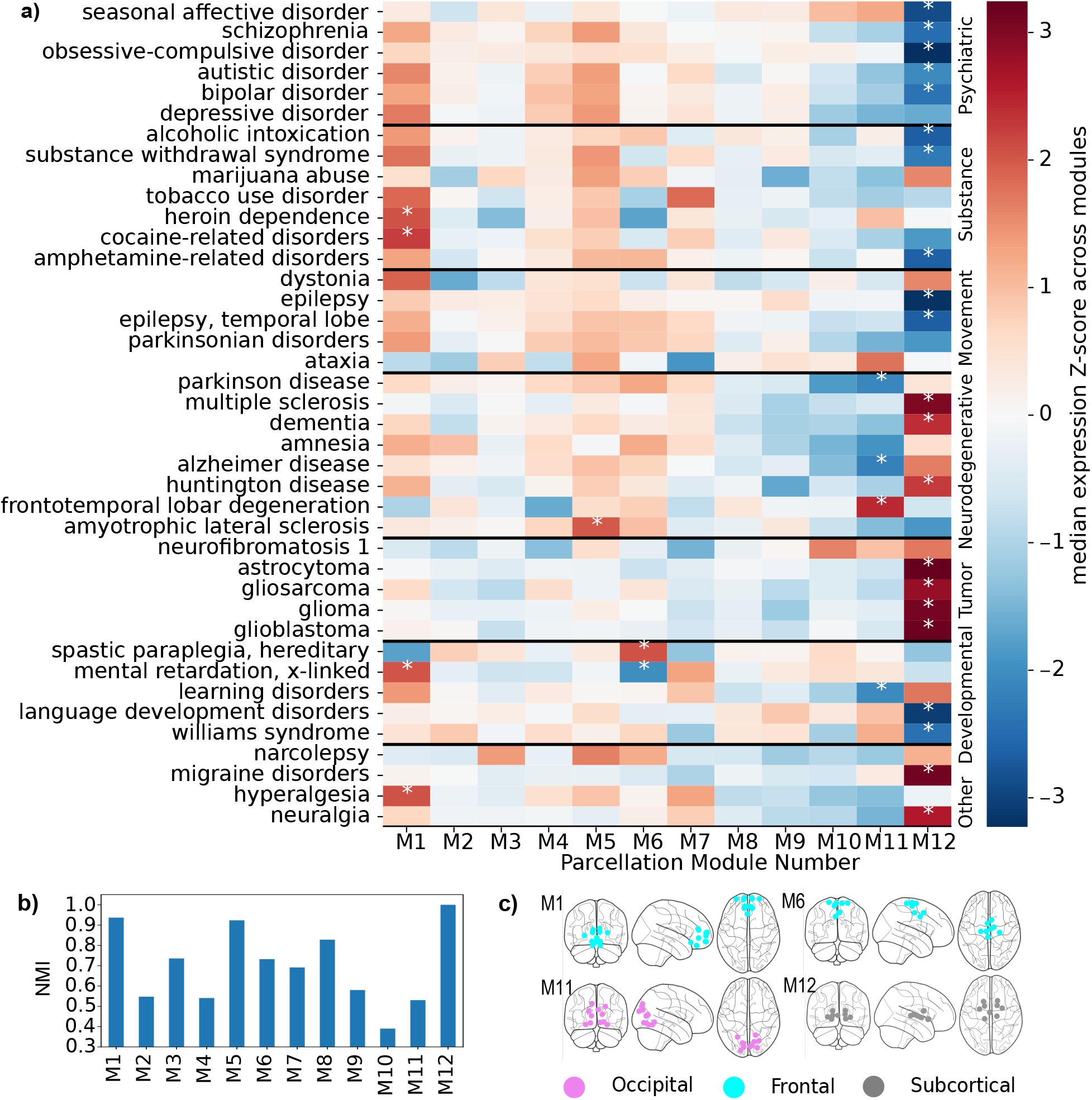
Transcriptomic expression of major brain-related disorders across different modules. For the modules of the same partition of the population 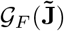illustrated in Fig. 2d, we show **a** the across-modules Z-score of the median representation of the genes associated with each disorder. Modules with absolute values higher than 2 (meaning that the disease is significantly over or under-represented) are denoted by a white asterisk. Histogram **b** contains the persistence of the partition’s modules across bootstrap samplings of the dataset’s subjects; the result is expressed in terms of the average match (NMI) of each module with its most similar counterpart in each sampling’s partition. Subfigure **c** shows the few modules significantly associated with three or more diseases. Notice the fundamental role of module M12, which is notably present across all samplings.

## 3 Discussion

In the study of brain structure-function coupling (SFC), functional connectivity (FC) inferred from partial correlation matrices shows a higher correspondence with structural connectivity (SC), even though most previous studies have only evaluated this correspondence at the single link level. In this work, we compared different methods to infer brain FC from resting-state BOLD time series. These methods include estimating FC from the correlation (**C**) or the partial correlations 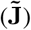 of the time series, inferred through a regularization algorithm, and further enhancing FC graph sparsity by setting the smallest (in absolute value) matrix elements to zero, up to the so-called percolation threshold.

In general, and in line with other works ([41, 42]), we found that inferring the FC graph from regularized partial correlation, rather than from correlation, enhances its sparsity and its similarity to the SC graph. While this is expected in the Gaussian, time-uncorrelated approximation, where the partial correlation represents the direct interactions between brain areas (also see S4 in SI), this result is by no means obvious in fMRI data, for several reasons. On the one hand, one expects fMRI data to be non-linear and time-correlated and, on the other hand, the estimations of the partial correlation-based FC are strongly influenced by the data finiteness and temporal correlations. Our article follows the above-cited studies, providing evidence of a closer FC-SC match in terms of partial correlations. Furthermore, we step further, by assessing: i) the importance of a noise-cleaning or regularisation method; ii) the impact of different regularisation methods; iii) the behaviour of a different metrics that accounts for the similarity between FC and SC at the hierarchical-modular level (beyond the element-wise covariance between the two matrices).

The utility of having a reliable, statistically significant estimator of connectivity, similar to SC, from BOLD time series has already been pointed out, at least in the context of computational approaches to brain function. There, the brain structure is often represented in terms of latent, interpretable, inferred parameters *θ*, that are eventually used as an input for machine learning classifiers, or to detect differences between different groups of subjects. Such a representation is called *generative embedding* ([64, 65, 66, 67, 68]). While a standard inference method of structural parameters *θ* from the imaging data is Dynamic Causal Modelling (DCM) ([69, 70]), a simpler, alternative to DCM is the linear scheme addressed here. Albeit linear models are much simpler and less realistic than DCM, their inference could be more statistically robust for a large number of brain areas and low number of time points, as is typical of fMRI time series.

As previously noted, retrieving FC from time series data requires statistical regularization. Here, we have addressed the robustness of the FC-SC match with respect to the statistical regularization method (Fig. 2), as well as with respect to the value regularization parameter, given the method. Rather interestingly, we find that the statistical regularization procedure, which has no information about SC, already brings the ensuing FC networks as similar to SC as those obtained by tuning the regularization parameter to the value *α*_*χ*_ that maximises the FC-SC match at the level of the single subject (Fig. 3).

In this work, we observed that working with sparse graphs is crucial for observing a high degree of cross-modularity. This is why we thresholded all graphs at the so-called *percolation threshold* (also see Methods) that depends on the graph topology, so that no arbitrary choice regarding the final density has to be made. Some authors have argued that thresholding is not a principled approach, as it results in a non-positive definite matrix, which does not represent a valid covariance matrix and may not be invertible, thus preventing association with a Gaussian likelihood ([41]). Consequently, they recommend using methods such as GLASSO to recover the adjacency matrix of 𝒢_*F*_ as a sparse partial correlation (a mathematically valid correlation matrix) whenever sparsity is required. Indeed, we confirmed that the GLASSO regularization, tuned by maximizing the test-set likelihood, produces a sparse PC matrix, whose cross-modularity is only slightly smaller than the one obtained by further thresholding the PC matrix, up to percolation value (Fig 4).

Given that the graphs’ density is highly relevant to the cross-modularity (as well as to many other graph properties ([63])), we analysed this dependency (see Fig. 4a) in order to rule out the hypothesis that the enhanced SC-FC match of 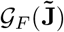 is only due to its higher sparsity, with respect to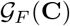. Our analysis reveals that, as a matter of fact, the higher SC-FC match for partial correlations occurs for a wide range of values of the density of FC connectivity matrices, and is not simply induced by the relatively low density of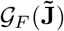. Fig. 4a also tells us that the lower the density, the higher the FC-SC match. Since the density is lowered by increasing the threshold *σ* below which we cut away the elements of the connectivity matrix, this fact suggest that higher similarity between FC and SC is obtained with the stronger links of the FC matrices.

Finally, our neurogenetics association analysis shows that Module 12, anatomically comprising the basal ganglia and thalamus -involved in motor control, cognition, and emotional regulation-, has a more dominant implication across different disease groups (Fig. 5). Increasing evidence suggests that gene expression patterns within the basal ganglia-thalamocortical circuitry overlap significantly with genetic profiles associated with various neuropsychiatric conditions, as well as mood and compulsive disorders. In depression, dysfunctions in the striatum and thalamus correlate with anhedonia and impaired reward processing, with genetic studies highlighting disrupted expression of serotonin transporters and dopaminergic genes ([71]). Similarly, obsessive-compulsive disorder is associated with hyperactive cortico-striato-thalamic loops, where altered expression of glutamatergic and serotonergic genes plays a role ([72]). In bipolar disorder, irregularities in the basal ganglia and thalamus contribute to emotional dysregulation ([73]). In autism spectrum disorders, disruptions in synaptic excitation-inhibition imbalance and neurodevelopmental gene expression in these regions contribute to sensory-motor dysfunction and social impairment ([74, 75]). Our neurogenetic results also show that both the basal ganglia and thalamus are involved in seizure modulation. Indeed, the thalamus, particularly the centromedian nucleus, plays a central role in seizure propagation, with gene expression studies implicating mutations in SCN1A and GABRG2 in epileptogenesis ([76]). On the other hand, the basal ganglia, particularly the substantia nigra pars reticulata, is involved in seizure suppression, and altered dopamine receptor gene expression has been observed in epilepsy models ([77]). In Alzheimer’s disease, the thalamus and basal ganglia exhibit significant atrophy, and recent transcriptomic analyses have shown that genes such as APOE, MAPT, and PSEN1 are differentially expressed in these regions ([78]). Huntington’s disease, which primarily affects the striatum, is characterized by mutant HTT gene expression, leading to neuronal loss and network dysfunction between the basal ganglia and thalamus ([79]).

## 4 Materials and Methods

### 4.1 Data

Connectivity data, downloaded from the open dataset ([53]), include the preprocessed resting-state functional magnetic resonance imaging (rs-fMRI) hemodynamic blood oxygen level-dependent (BOLD) time series and the Diffusion Weighted Imaging (DWI) structural connectivity matrices of 136 healthy participants, 98 of them males, aged between 20 and 30 years old, already parcelled in 183 regions of interest (ROIs). The rs-time series include 652 time steps. Details about the preprocessing and the parcellation, from raw data of the multimodal dataset *Max Planck Institut Leipzig Mind-Brain-Body Dataset* commonly referred to as LEMON ([80]), can be found in ([31]).

Transcriptomic expression data, containing the expression of genes within each ROI, were also obtained from the dataset ([31]), and they are the result of the preprocessing, using the *abagen* tool ([81]), of the open data from the Allen Human Brain Atlas (AHBA) ([82]). In addition, we obtained the data relative to the association between genes and diseases from the archive associated with ([54]).

### 4.2 From data to brain Functional and Structural graphs

The SC and FC graphs were recovered from the respective connectivity matrices, interpreted as adjacency matrices: each matrix entrance can be read as the strength of the connection (*link weight*) between the corresponding couple of ROIs (*nodes*). For each subject *s*, we obtained the SC directly from the structural data, while we computed multiple estimates of the FC’s connectivity matrix from the resting state BOLD time series: from their regularized correlation matrix 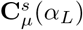 and from its inverse 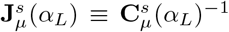, whose elements were next normalized as 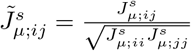 (*partial correlation*), for each of the regularization methods *µ* that we took into account (see next section); the regularization parameter *α*_*L*_ maximes the likelihood of the validation set, with respect to the method. The “raw,” not regularized, correlation matrix is the sample covariance of the fMRI time series; it corresponds to the sample correlation matrix when the time series are demeaned and standardized (null temporal averages and unit standard deviation) as we assume the data to be. Since there is no straightforward interpretation for negative link weights, the adjacency matrices **M** of the FC connectivity graphs were taken as the **C**, 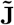 matrices in absolute value. More specifically, 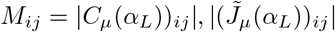, for the correlation-based and for the partial correlation-based FC networks (i.e., 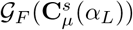 and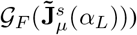, respectively.

Next, we cut both structural and functional matrices **M** at the *percolation threshold* (for more detail see the dedicated section). In the preliminary analysis, we verified that the way negative values are treated does not significantly affect the results since most of the negative elements are small in absolute value and are consequently removed in the thresholding step. At last, the thresholded matrices **M** are taken as the adjacency matrices of the SC and FC graphs of each subject *s*.

At the population level, we defined the population connectivity matrix as the across-subject median of the single-subject connectivity matrices.

### 4.3 Regularization methods

Given the single subject time series **X** (a *N* × *T* real matrix), demeaned and standardized, if *T* is finite or *q* = *N/T* does not vanish, the sample correlation matrix 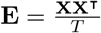 and, even more, its inverse **J** = **E**^*−*1^ are not good estimators of the true correlation and precision matrices ([48, 62, 45]). Regularizing, or noise-cleaning, consists of proposing a matrix that is as similar as possible to the true correlation matrix (in particular, more than **E**).

The regularization algorithms that we took into account in this work (far from being a complete list of all the methods present in literature ([62])), include methods from the two most common approaches in neuroscience (i.e., the Linear Shrinkage approach, represented by the Identity and Group Shrinkage, and the sparse estimator approach, such as Graphical Lasso ([41])), plus two algorithms that follow a Principal Component approach. All these algorithms were systematically evaluated in ([45]) on rs-fMRI BOLD signal and synthetic data, in terms of scores such as the element-wise distance *d*(**J, J**^*v*^) between the inferred precision **J** and the true precision **J**^*v*^, and the test-set likelihood. Most of the regularization algorithms consist in regularizing the spectrum **Λ** of **E** while keeping the eigenvectors **U** unchanged (based on the assumption of no prior knowledge of the eigenvectors’ structure) so that the correlation matrix regularized with method *µ* has the general form **C**_*µ*_(*α*) = **UΛ**(*α*)**U**^⊺^: a,b) **Linear Shrinkage (LS)** ([83]), consists in a convex combination **C**_*LS*_(*a*) = (1 − *a*)**E** + *a***T** between the sample matrix **E** and a target matrix **T** (independent of the data), such as the identity matrix 𝕀_*N*_ (in which case we dub the method **Identity Shrinkage (IS)**), or the across-subject average of the sample correlation matrix *⟨***E***⟩* (we call this method **Group Shrinkage (GS)**). The Linear Shrinkage can have different interpretations. For example, it can be seen as a trade-off between bias and variance, as a shrinkage of the eigenvalues towards their grand mean ([83]), or also as an optimal Bayes estimator in the context of the Bayesian Random Matrix theory, choosing the minimum mean squared error as loss function and assuming a Gaussian distribution for the data and an Inverse-Wishart as prior distribution for the correlation matrix ([62]).

**c) Principal Component Analysis (PCA)**, (also known as *Eigenvalues Clipping* ([62])) that consists in considering as significant only the *p* largest eigenvalues of **E**: 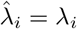if *i ≤ p*, else 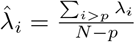.

**d) Cautious Principal Component Analysis (cautPCA)**, a simple variant of PCA that was firstly introduced in ([45]), where it proved to slightly outperform PCA. In this case the spectrum is first changed as 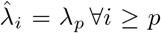and later rescaled such that *tr* [**C**] = *N*.

**e) Graphical Lasso (GLASSO)**; in this case **C** and **J** are computed by maximization of the log-likelihood minus a *ℓ*_1_ norm penalty term: **J**_*GLASSO*_(*a*) = *arg* max_*J*_ ln {*N* (**X** | **J**^*−*1^) −*a ∑* _*i<j*_ | *J*_*ij*_|} ([46])

**f) Optimally Rotationally Invariant Estimator (ORIE)**; first proposed in ([84]) and later extended in ([85]), the method consists of correcting the eigenvalues of the sample estimator **E** as 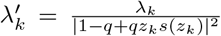 where *s*(*z*) = *Tr*[(*z*𝕀_*N*_ − **E**)^*−*1^]*/N*, and *z*_*k*_ = *λ*_*k*_ − *iη* (*i* is the imaginary unit) ([84, 85, 86, 62, 45]). This estimator minimizes, among all estimators that share the same eigenvectors of the sample correlation, the Euclidean distance from the true correlation matrix in the high-dimensional limit. The parameter *η* should be small, and such that *Nη ≫* 1 ([86]). While a convenient choice is given by *η* = *N*^*−*1*/*2^ ([86, 85, 45]), this may lead to values that are not small enough if *N* is not extremely large ([86]). Therefore, following ([45]), we choose to cross-validate this parameter to maximize the validation-set likelihood.

**Figure S1:**
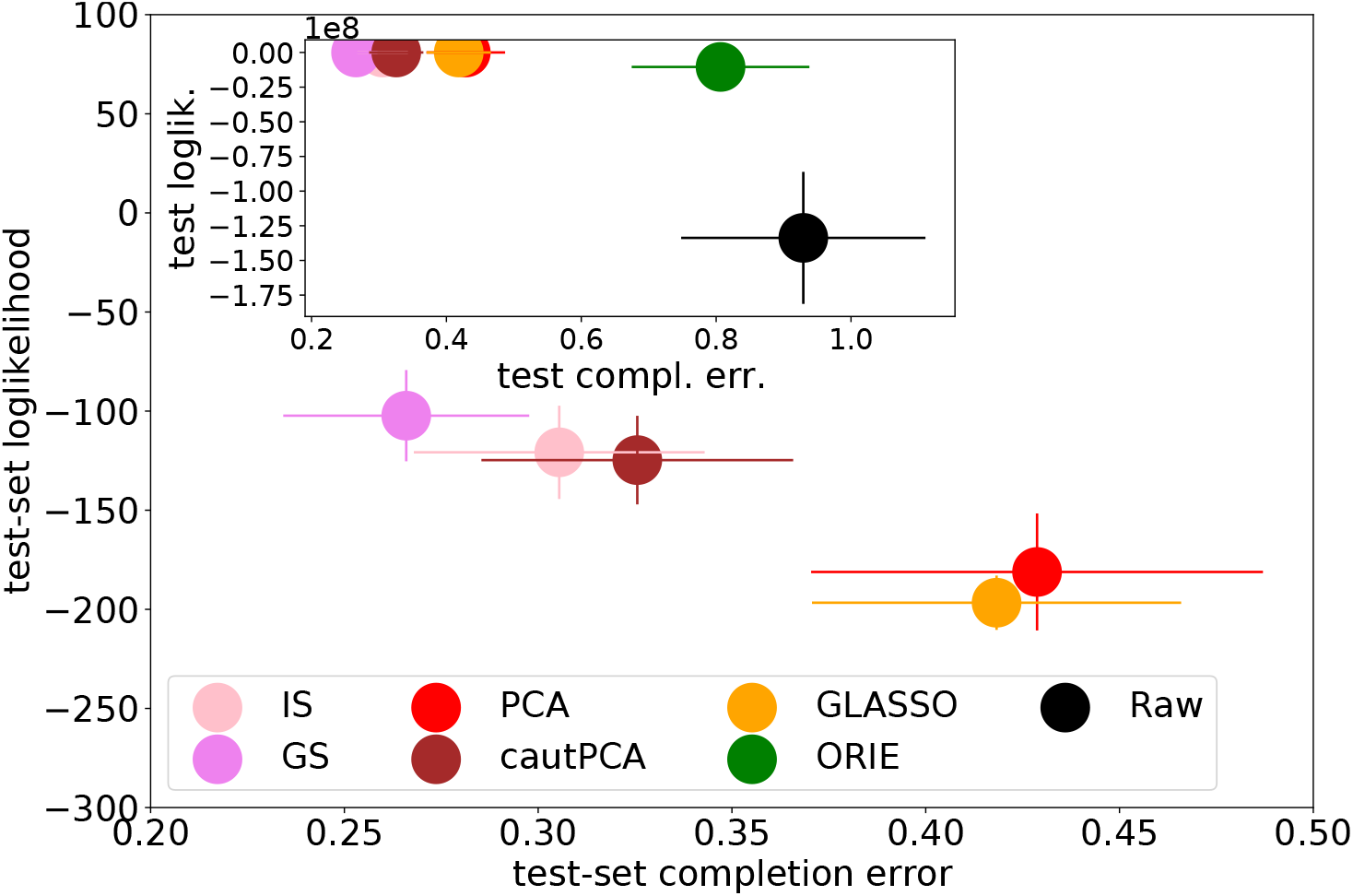
Values of the test-set likelihood. Scores of the regularized correlation (and hence precision) matrices, computed on the training set, in terms of test-set likelihood and completion error (see [45]), for different regularization methods. “Raw” stands for no regularization. Importantly, for each subject, the training and test sets are obtained randomly splitting the time series (time-wise), without altering the temporal order of the vectors.

Except for ORIE and GLASSO, the regularization methods that we took into account in this work depend on a tuning parameter that we generically call *α*, ranging from zero (corresponding to no regularization at all, in which case the correlation matrix estimate equates the sample correlation **E**) to *α* = 1 (maximum regularization). The regularization parameter coincides with the parameter *a* of LS and with 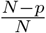 for PCA and cautPCA.

As explained in more detail in SI, S1, the regularization parameters are set, by cross-validation, to those maximizing the validation-set likelihood at the level of the single subject BOLD time series.

### 4.4 Percolation threshold

As mentioned, we obtained the connectivity graphs from the respective thresholded connectivity matrices. In particular, we thresholded the matrices at the percolation cutoff, meaning that we gradually removed the matrix elements in order of increasing weight, stopping just before the corresponding graph would break into more than one disconnected component. Among the many sparsification methods that have been proposed in neuroscience, this method has the advantage of providing a connected graph whose final density depends on the graph topology. Moreover, in ([55]), the application of the percolation threshold was shown to provide the optimal balance between the removal of noise and genuine information, maximizing the distance of FC from its randomized counterpart, and it was suggested that its application is critical for the extraction of the large-scale structure from the network.

### 4.5 Hierarchical Clustering

We compared the architectures of each pair of SC and FC graphs, both at the subject and population levels, based on their hierarchical clusterings. Hierarchical, or agglomerative, clustering is an unsupervised method for finding communities in N-dimensional observation vectors. The algorithm produces a whole hierarchy of nested data partitions that can be represented as a dendrogram. As a result of this step, we obtained a couple of sets of partitions 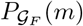 and 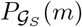 in *m* modules, *∀m* = 2, …50, representing the hierarchical partitions of both Functional and Structural Connectivity graphs.

In particular, we have made use of a hierarchical clustering algorithm whose distance matrix is built as exp(−**M**_*ij*_), where **M** is the connectivity graph adjacency matrix. More in detail, we used SciPy implementation module ([87]), with `metric=cosine` metric and `method=weighted`, taking as input the so called *1-D condensed distance matrix* built from the adjacency matrix.

### 4.6 Cross-Modularity

To evaluate the similarity of the partitions of the Functional and Structural Connectivity, we introduced a metric that, given the partitions 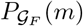 and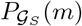, simultaneously takes into account their individual quality and their reciprocal similarity, for each number of modules *m*. Therefore we define the cross-modularity *χ*(*m*) as

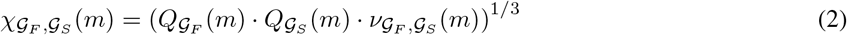

where *Q*𝒢 (*m*) is the quality of partition *P*𝒢 (*m*), measured in terms of the Newmann Modularity ([56]) (we measured it ignoring the links’ weights), and 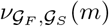is the similarity of partitions 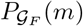 and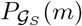, measured in terms of Adjusted Normalized Mutual Information (NMI) ([57]). The cross-modularity is a variation of the homonym metric first introduced in ([30]); there, a unique partition *P* (*m*) was applied to both SC and FC graphs, and the agreement between the communities of the two graphs was then computed as the average agreement (measured as Sorensen index) between the couple of communities induced on SC and FC by each module of *P* (*m*).

More in detail, Newman modularity is a measure of the quality of a particular partition of a graph into modules, defined in ([56]) as:

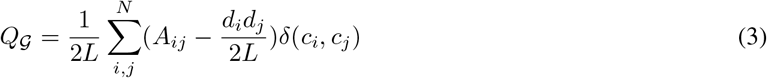

where *L* is the number of edges, *A* the adjacency matrix, *d*_*i*_ the degree of *i* and *δ*(*c*_*i*_, *c*_*j*_) is 1 if *i* and *j* belong to the same community, else 0. This quantity is proportional to the number of intra-module edges minus its expected number in a network with equal degree sequence and random links. Consequently, modularity approaching 1 indicates a high-quality partition (high intra-cluster and low inter-cluster density), while it would vanish in the case of a random one ([59]).

Mutual Information (MI) is an information-theoretic tool that measures the amount of information shared by two partitions; its normalized and adjusted version (NMI), where the adjustment discards the matches due to chance, was proposed in ([61]):

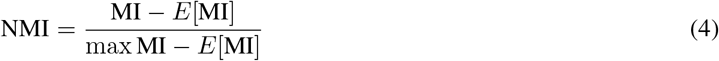

(where symbol *E*[·] stands for the expected value), so that this quantity vanishes in case of comparison of random partitions and reaches 1 for perfect coherence.

As a consequence, cross-modularity may achieve a maximum value of 1 in the case of high-quality partitions and a perfect match, while it vanishes in the case of unrelated partitions or if at least one of them is random. We have measured Newman modularity using the implementation in ([58]) and NMI using the implementation in ([60]).

### 4.7 Spectral distance

In addition to the single-subject cross-modularity, we also computed the spectral distance ([88]) of the connectivity graphs’ adjacency matrices between each couple of subjects for both the SC graph and every estimate of the FC graph.

In general, the spectral distance measures the difference between a couple of matrices in terms of the difference between their eigenvalues:

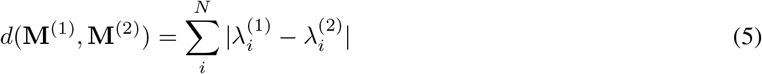

where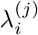, with *j* = 1, 2 and *i* = 1, …, *N* are the matrices **M**^(1)^ and **M**^(2)^ eigenvalues. To make fair the comparison between the spectral distance of the subjects’ FC (which are bounded, in absolute value, between 0 and 1) and SC (which are expressed as the number of white matter streamlines) adjacency matrices, we normalized the values of the (thresholded) 𝒢_*S*_ adjacency matrix as 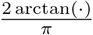.

### 4.8 Characterization of the optimal FC brain partition in terms of brain-related disorders

Once obtained the optimal (in the sense of higher cross-modularity) partition of the optimal (in the sense of statistical regularization) estimate of the FC graph at the population level, we characterized its modules depending on their participation in 40 major brain-related disorders from 7 disease groups (Psychiatric disorders, Substance abuse, Movement disorders, Neurodegenerative diseases, Tumor conditions, Developmental disorders, and Others). Our method follows the procedure adopted in ([31]), work associated with the Zenodo dataset from which we extracted the transcriptomic data. For each disease and for each module, we computed the disease expression value as the median value across the genes associated with the disease and across the nodes contained in the module. The heatmap in Fig. 5 shows the across-modules Z-scores of the diseases expressions, with absolute values higher than 2 marked with an asterisk, and diseased grouped by the WHO categories (also obtained from archive ([54])).

### 4.9 Robustness check of the optimal FC brain partition

We have checked the robustness of the partition of our estimate of the population FC graph by comparing it with different estimates inferred from 100 bootstrap samplings of the dataset. In particular, for 100 times we iterated the following procedure: we randomly selected 136 subjects (importantly, with repetitions), we computed an estimate of the population 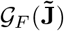 of these subjects, with regularization method GS and we partitioned it into 12 modules; finally, we computed the match of each partition with the whole-dataset one (i.e., the partition shown in Fig 2) in terms of NMI, finding an overall score of 0.86. We additionally measured the robustness of each whole-dataset partition’s module as its across-sampling average NMI similarity to the most similar module in the subsampling partition. The results, shown in Fig. 5b, reveal that some modules, module M12 *in primis*, are particularly robust.

## 5 Acknowledgments

This work has been supported by: the European Union under the scheme HORIZON-INFRA-2021-DEV-02-01 - Preparatory phase of new ESFRI research infrastructure projects, Grant Agreement n.101079043, “SoBigData RI PPP: SoBigData RI Preparatory Phase Project”; by the project “Reconstruction, Resilience and Recovery of Socio-Economic Networks” RECON-NET EP_FAIR_005 - PE0000013 “FAIR” - PNRR M4C2 Investment 1.3, financed by the European Union - NextGenerationEU; by the European Union - Horizon 2020 Program under the scheme ‘INFRAIA-01-2018-2019 - Integrating Activities for Advanced Communities’, Grant Agreement n.871042, ‘SoBigData++: European Integrated Infrastructure for Social Mining and Big Data Analytics’ (http://www.sobigdata.eu); Grant No. PID2023-149174NB-I00 financed by MICIU/AEI/10.13039/501100011033 and EDRF/EU funds.

Jesus M. Cortes acknowledges financial support from Ikerbasque: The Basque Foundation for Science, and from Spanish Ministry of Science (PID2023-148012OB-I00), Spanish Ministry of Health (PI22/01118), Basque Ministry of Health (2023111002 & 2022111031).

## Supplementary Information

### S1 Statistical Regularization

The regularization methods that we took into account rely on a tuning parameter *α*. Its optimal value *α*_*L*_ is set by maximization of the log-likelihood ln (*P* (**X**_*v*_ | **C**_*tr*_)) evaluated on the *validation-set* **X**_*v*_, different from the *pure-training-set* **X**_*tr*_ used to infer **C**_*tr*_ (we adopt the notation of reference ([45]), in which **X** = ((**X**_*tr*_, **X**_*v*_), **X**_*te*_), where the joint pure-training and validation-sets (**X**_*tr*_, **X**_*v*_) constitute the training-set, and where “te” refers to test-set). We perform the maximization of the validation-set likelihood across a grid of values of *α* through k-fold cross-validation ([60]). The training set (**X**_*tr*_, **X**_*v*_) is randomly split into *k* smaller sets, or folds; at each of the *k* iterations, *k* − 1 folds are taken as pure-training set (**X**_*tr*_), and the remaining one as validation-set (**X**_*v*_); finally, each parameter’s score is computed as the average of the iteration scores. In Fig. S1 we show the test-set likelihood of the main dataset corresponding to each regularization method. The statistical regularization methods were implemented using the repository ([89]).

An important detail regarding the cross-validation procedure is that we respect the original temporal order of the series (the index *t* of *X*_*jt*_) in each fold of the pure-training-set/validation-set division. We use, in other terms, the flag `shuffle=False` of the function `split_train_test` of ([89]). In the presence of a high correlation time (as the one exhibited by the dataset analysed in the main text), we notice that it is necessary not to shuffle the temporal order of the data in the maximization of the validation-set likelihood by cross-validation. In short, if the correlation time of the data is of the same order of the number of samples in the validation-set, and if one shuffles the temporal order, the validation-set likelihood can be strongly overestimated for low values of the regularization parameter, since the validation-set vectors are too similar to some vectors in the pure-training-set. As a consequence, the consequent values of *α*_*L*_ may be severely underestimated. As a consistency check of these arguments, we have verified that in the supplementary extra dataset (see Fig. S2), which exhibits a significantly lower correlation time, shuffling or not shuffling the temporal order leads to qualitatively identical results (in their turn, consistent with the results found for our main dataset, without shuffling), also in terms of FC-SC cross-modularity. This point will be explained in detail in an upcoming publication.

**Figure S2:**
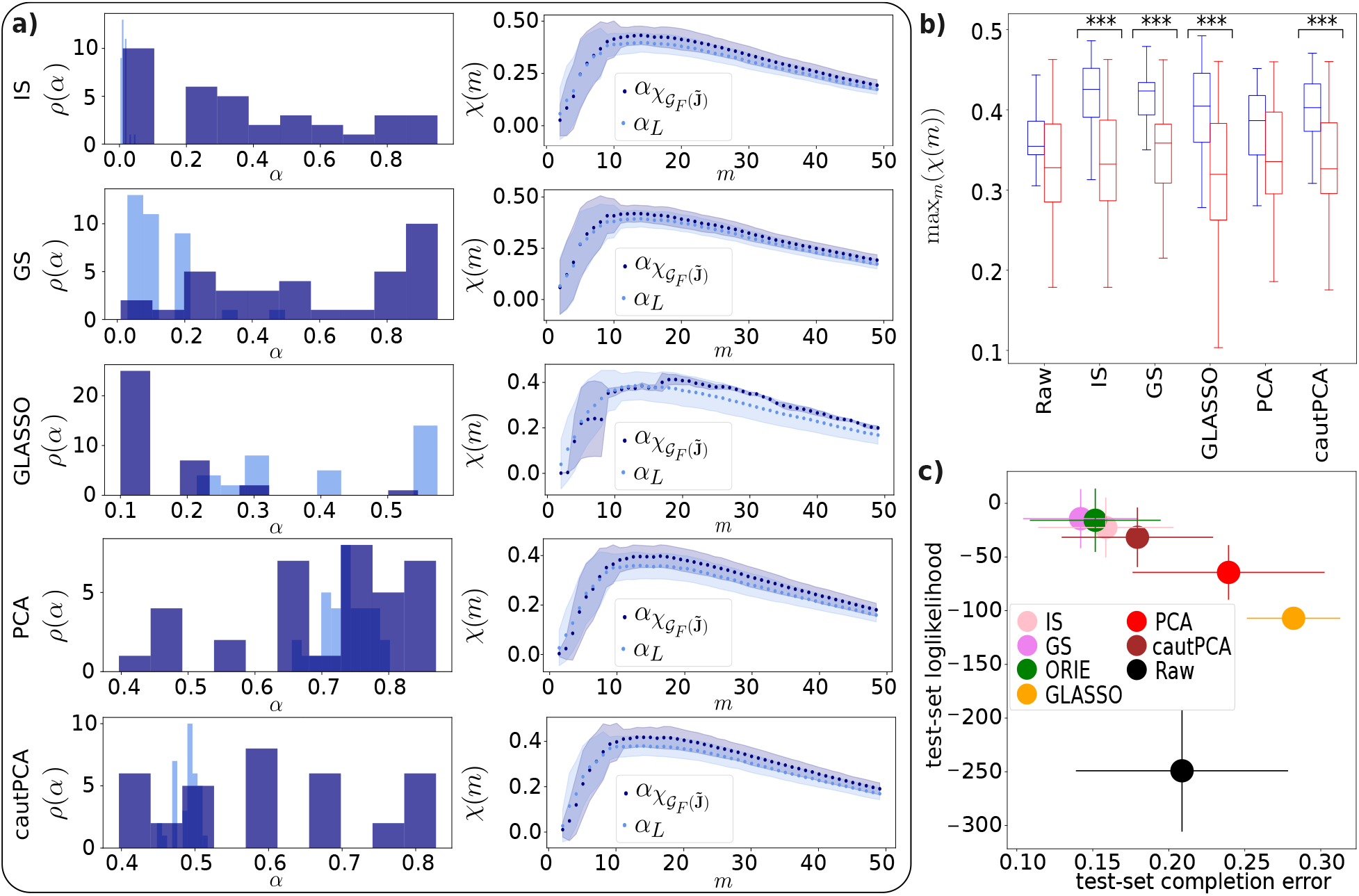
Regularization tuning parameters. (subfigure **a**) and maximum cross-modularity (subfigure **b**) computed on a **supplementary extra dataset**, and **test-set likelihood scores** (subfigure **c**). The extra dataset is composed of 35 subjects, with 116 ROIs and rs-BOLD time series of 180 time steps. Coherently with our main results (see Fig. 3), we do not observe any significant gap between the cross-modularity of 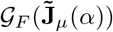 choosing the tuning parameter *α*_*L*_ that maximizes the validation-set likelihood and 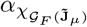 that maximizes the cross-modularity of 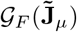(subfig. **a**). In addition, the subjects’ *χ*(*m*) curves exhibit a maximum at approximately the same value of *m* as in the main dataset (subfig. **a**). As in the case of the main dataset, the cross-modularity of the FC graph retrieved from regularized partial correlation is significantly higher than that retrieved from correlation (subfig. **b**; also compare with Fig. 2. The three asterisks indicate the methods corresponding to a p-value lower than 10^*−*3^ for the Mann-Whitney U rank test of the null hypothesis that the distribution underlying the sample corresponding to **C** is stochastically less than the distribution underlying the sample corresponding to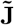).

### S2 Consistency analysis for the supplementary dataset

We performed a consistency analysis of the main article results in a supplementary dataset. The supplementary dataset consists the rs-fMRI BOLD series and DWI matrices of 35 subjects, with *N* = 116 ROIs, and *T* = 180 time steps (please, see the details in ([90]). Fig. S2 summarizes our results (compare with Figs. 2, 3). We observe that the article results are perfectly consistent across datasets.

### S3 FC-SC correspondence at the population level

We show here in detail the effect of the regularization on *χ*(*m*) at the population level, having selected the simple GS as reference method. Figs. S3a-c provide a representation of the population 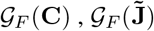 and 𝒢_*S*_. From a visual inspection, 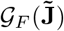 appears characterized by a higher modularity than 𝒢_*F*_ (**C**) and by a lower density, closer to that of 𝒢_*S*_.

Figs. S3d,e reveal that 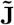 provides a higher cross-modularity than **C**, but the effect of regularization is less significant compared to the single-subject case.

**Figure S3:**
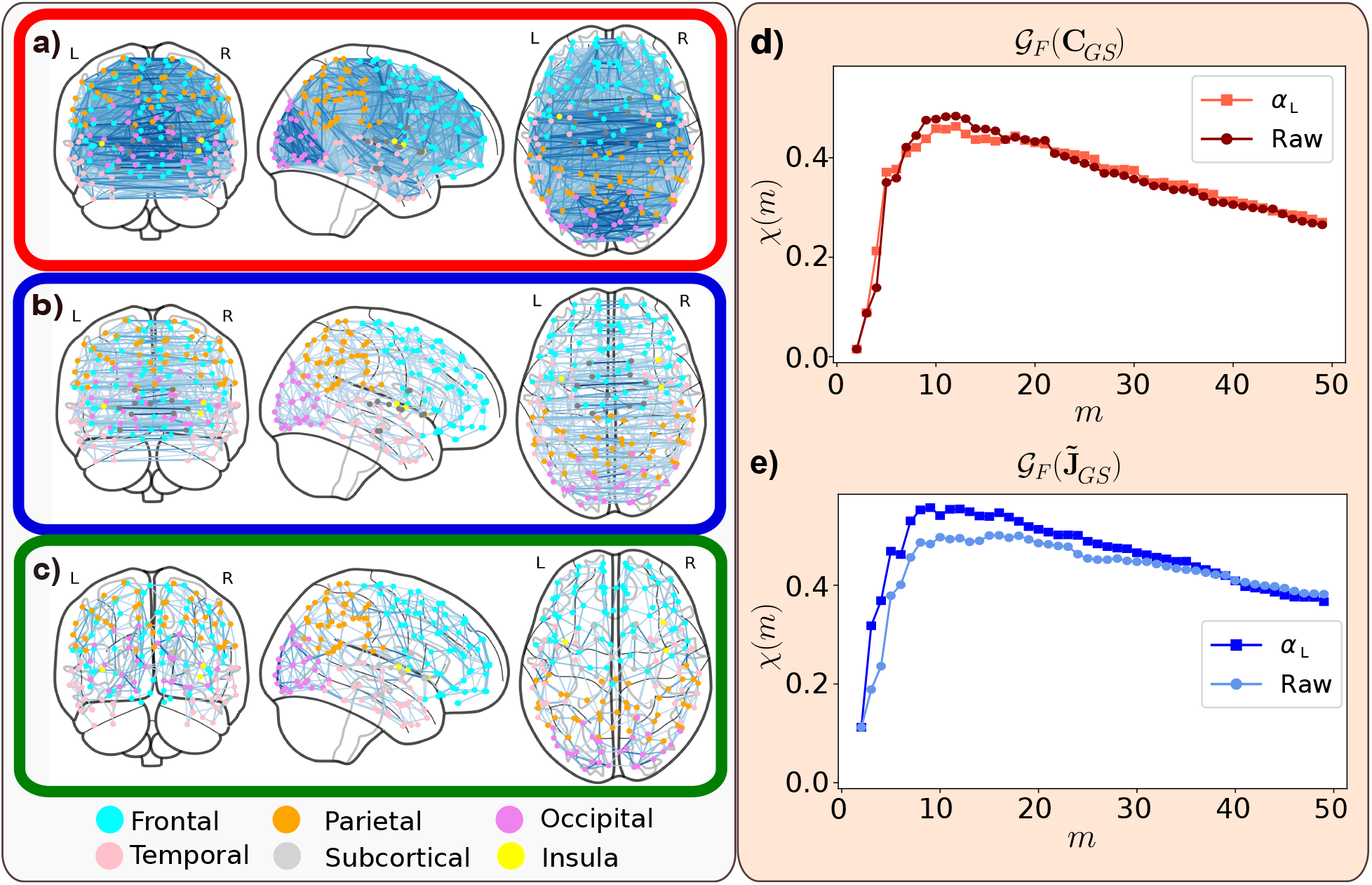
Connectivity graphs, inferred from C and 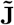, at the population level. The first column reveals the aspect of the population brain graphs, where darker edges correspond to stronger links: 𝒢_*F*_ (**C**) (**a**, red frame), 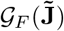(**b**, blue frame) (both computed as the median of the single subject matrices regularized with GS), and 𝒢_*S*_ (**c**, green frame). A visual inspection appears in agreement with the rest of the results, suggesting that 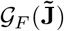 is closer to G_*S*_ than G_*F*_ (**C**). The second column shows the population cross-modularity *χ*(*m*), for 𝒢_*F*_ (**C**) (**d**, red) and 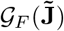 (**e**, blue), for not regularized (“Raw”) and regularized (*α*_*L*_) matrices.

### S4 Relation between Structure and Function in the linear approximation

In this section, we provide a detailed discussion about why we expect partial correlations to be closer (than simple correlations) to structure, and what are the working hypothesis underpinning this expectation.

The working hypothesis of this and other articles considering full and partial covariance matrices is a linear interaction between different brain areas (see, for example, [52, 40, 42]). In other words, the hypothesis is that the time series’ stationary probability distribution is normally distributed with a certain correlation matrix **C** (and associated precision matrix **J** = **C**^*−*1^), to be inferred:

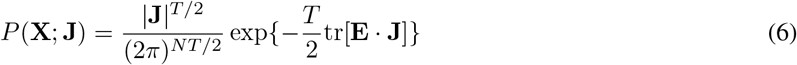

where (we assume the time series **X** to be demeaned and standardized)

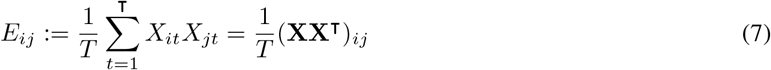

is the sample covariance, or, equivalently,

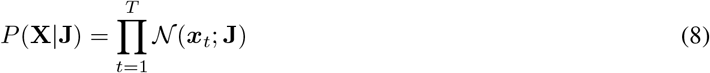

where

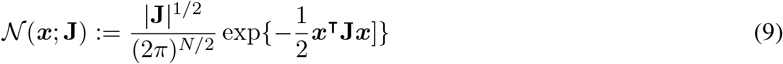

is the multivariate normal distribution and where

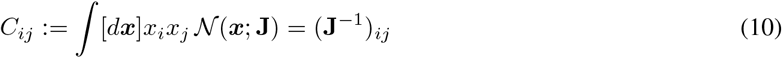

**Figure S4:**
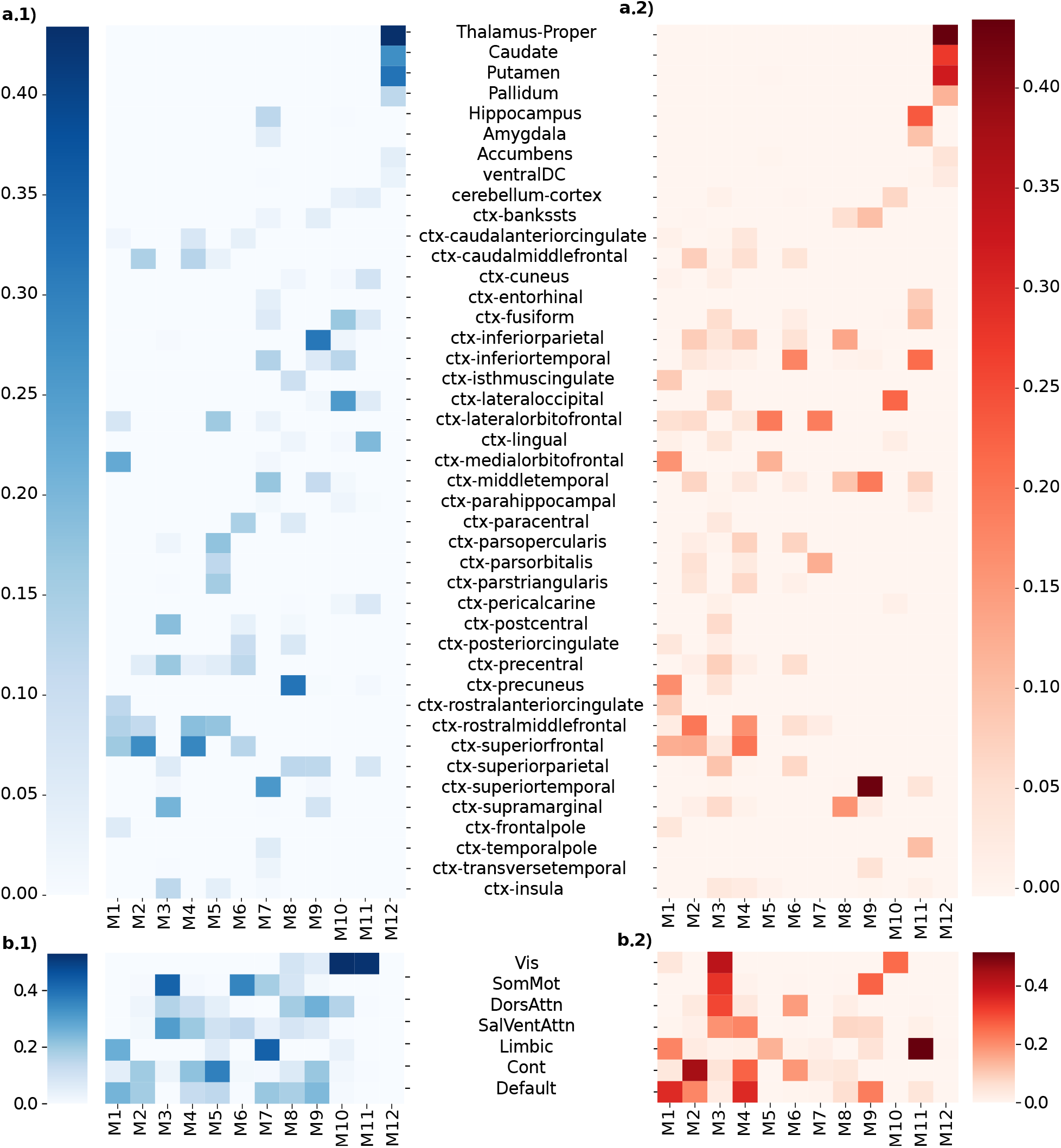
Overlap between modules and brain areas. This image shows the overlap between each module of both the population 𝒢_*F*_ (**C**) (red) and 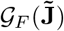 (blue), regularized with GS (in the case of 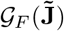 the modules are the same as those shown in Fig. 2), the Desikan-Killany structural areas (subfigures *a*), and the Yeo functional areas (subfigures *b*). The overlap is measured in terms of the Jaccard index between each couple module-area’s volumes.

This assumption provides a probabilistic relation *P* (**X** | **J**) between different areas’ time series **X**, induced by the precision matrix **J** that, in a statistical physics language, represents the (harmonic) interactions between couples of brain areas. More in detail, *x*_*i*_ and *x*_*j*_ represent the deviations of areas *i* and *j* from their respective baseline mean BOLD activities, coupled by causal, harmonic interactions (i.e., by a virtual spring with elastic constant *J*_*ij*_, to be inferred) that induce an elastic interaction energy equal to *ϵ*_*ij*_ = *x*_*i*_*x*_*j*_*J*_*ij*_. Thus, *P* (**X** | **J**) corresponds to the Bolztmann probability distribution associated with the energy induced by the “structural” constraints **J**. Such an interaction matrix induces, in turn, the covariances **C**. Therefore, **J** represents the direct, causal interactions, while **C** represents the emerging (direct and spurious) covariances, associated with a probability distribution *P* (**X** | **J**) over the ensemble of all possible configurations of BOLD activity. Therefore, in a linear approximation, the relation between structural connections **J** and associated, emerging covariances **C** is simply **C** = **J**^*−*1^.

## References

[1] Karl J Friston. Functional and Effective Connectivity in Neuroimaging: A Synthesis. Human Brain Mapping, 2(1-2):56–78, 1994.

[2] Christopher J Honey, Rolf Kötter, Michael Breakspear, and Olaf Sporns. Network structure of cerebral cortex shapes functional connectivity on multiple time scales. Proceedings of the National Academy of Sciences, 104(24):10240–10245, 2007.

[3] Martijn P. Van Den Heuvel, René C.W. Mandl, René S. Kahn, and Hilleke E. Hulshoff Pol. Functionally linked resting-state networks reflect the underlying structural connectivity architecture of the human brain. Human Brain Mapping, 30(10):3127–3141, October 2009.

[4] Michael D. Greicius, Kaustubh Supekar, Vinod Menon, and Robert F. Dougherty. Resting-state functional connectivity reflects structural connectivity in the default mode network. Cerebral Cortex, 19(1):72–78, January 2009.

[5] CJ Honey, O Sporns, L Cammoun, X Gigandet, JP Thiran, R Meuli, and P Hagmann. Predicting human resting-state functional connectivity from structural connectivity. PNAS, 2009.

[6] Christopher J. Honey, Jean Philippe Thivierge, and Olaf Sporns. Can structure predict function in the human brain?, September 2010. ISSN: 10538119 Issue: 3 Pages: 766–776 Publication Title: NeuroImage Volume: 52.

[7] Hae-Jeong Park and Karl Friston. Structural and Functional Brain Networks: From Connections to Cognition.

[8] Joaquin Goni, Martijn P. Van Den Heuvel, Andrea Avena-Koenigsberger, Nieves Velez De Mendizabal, Richard F. Betzel, Alessandra Griffa, Patric Hagmann, Bernat Corominas-Murtra, Jean Philippe Thiran, and Olaf Sporns. Resting-brain functional connectivity predicted by analytic measures of network communication. Proceedings of the National Academy of Sciences of the United States of America, 111(2):833–838, 2014.

[9] Olaf Sporns and Richard F. Betzel. Modular brain networks. Annual Review of Psychology, 67:613–640, January 2016. Publisher: Annual Reviews Inc.

[10] Kanika Bansal, Johan Nakuci, and Sarah Feldt Muldoon. Personalized brain network models for assessing structure–function relationships, October 2018. ISSN: 18736882 Pages: 42–47 Publication Title: Current Opinion in Neurobiology Volume: 52.

[11] Bertha Vázquez-Rodríguez, Laura E. Suárez, Ross D. Markello, Golia Shafiei, Casey Paquola, Patric Hagmann, Martijn P. Van Den Heuvel, Boris C. Bernhardt, R. Nathan Spreng, and Bratislav Misic. Gradients of struc-ture–function tethering across neocortex. Proceedings of the National Academy of Sciences of the United States of America, 116(42):21219–21227, October 2019. Publisher: National Academy of Sciences.

[12] Laura E. Suárez, Ross D. Markello, Richard F. Betzel, and Bratislav Misic. Linking Structure and Function in Macroscale Brain Networks. Trends in Cognitive Sciences, 24(4):302–315, April 2020.

[13] T. Sarwar, Y. Tian, B. T.T. Yeo, K. Ramamohanarao, and A. Zalesky. Structure-function coupling in the human connectome: A machine learning approach. NeuroImage, 226, February 2021. Publisher: Academic Press Inc.

[14] Farnaz Zamani Esfahlani, Joshua Faskowitz, Jonah Slack, Bratislav Mišić, and Richard F. Betzel. Local structurefunction relationships in human brain networks across the lifespan. Nature Communications, 13(1), December 2022. Publisher: Nature Research.

[15] Gustavo Deco, Viktor Jirsa,R R Mcintosh, Olaf Sporns, and Rolf Kö. Key role of coupling, delay, and noise in resting brain fluctuations. Technical report, 2024.

[16] Carmen Alonso-Montes, Ibai Diez, Lakhdar Remaki, Iñaki Escudero, Beatriz Mateos, Yves Rosseel, Daniele Marinazzo, Sebastiano Stramaglia, and Jesus M. Cortes. Lagged and instantaneous dynamical influences related to brain structural connectivity. Frontiers in Psychology, 6, July 2015. Publisher: Frontiers Media S.A.

[17] Amor T. A, Russo R, Diez I, Bharath P, Zirovich M, Stramaglia S, Cortes J. M, de Arcangelis L, and Chialvo D. R. Extreme brain events: Higher-order statistics of brain resting activity and its relation with structural connectivity. Europhysics Letters, 111(6), October 2015.

[18] S. Stramaglia, M. Pellicoro, L. Angelini, E. Amico, H. Aerts, J. M. Cortés, S. Laureys, and D. Marinazzo. Ising model with conserved magnetization on the human connectome: Implications on the relation structure-function in wakefulness and anesthesia. Chaos, 27(4), April 2017. Publisher: American Institute of Physics Inc.

[19] P. Hagmann, O. Sporns, N. Madan, L. Cammoun, R. Pienaar, V. J. Wedeen, R. Meuli, J. P. Thiran, and P. E. Grant. White matter maturation reshapes structural connectivity in the late developing human brain. Proceedings of the National Academy of Sciences of the United States of America, 107(44):19067–19072, November 2010. Publisher: National Academy of Sciences.

[20] Karla Batista-García-Ramó and Caridad Ivette Fernández-Verdecia. What we know about the brain structure-function relationship, April 2018. ISSN: 2076328X Issue: 4 Publication Title: Behavioral Sciences Volume: 8.

[21] Panagiotis Fotiadis, Linden Parkes, Kathryn A. Davis, Theodore D. Satterthwaite, Russell T. Shinohara, and Dani S. Bassett. Structure–function coupling in macroscale human brain networks, October 2024. ISSN: 14710048 Publication Title: Nature Reviews Neuroscience.

[22] Alessandra Griffa, Enrico Amico, Raphaël Liégeois, Dimitri Van De Ville, and Maria Giulia Preti. Brain structure-function coupling provides signatures for task decoding and individual fingerprinting. NeuroImage, 250, April 2022. Publisher: Academic Press Inc.

[23] Zijin Gu, Keith Wakefield Jamison, Mert Rory Sabuncu, and Amy Kuceyeski. Heritability and interindividual variability of regional structure-function coupling. Nature Communications, 12(1), December 2021. Publisher: Nature Research.

[24] Maria Giulia Preti and Dimitri Van De Ville. Decoupling of brain function from structure reveals regional behavioral specialization in humans. Nature Communications, 10(1), December 2019. Publisher: Nature Publishing Group.

[25] Rui Cao, Xin Wang, Yuan Gao, Ting Li, Hui Zhang, Waqar Hussain, Yunyan Xie, Jing Wang, Bin Wang, and Jie Xiang. Abnormal Anatomical Rich-Club Organization and Structural–Functional Coupling in Mild Cognitive Impairment and Alzheimer’s Disease. Frontiers in Neurology, 11, February 2020. Publisher: Frontiers Media S.A.

[26] Jonathan Tay, Marco Düring, Esther M.C. van Leijsen, Mayra I. Bergkamp, David G. Norris, Frank Erik de Leeuw, Hugh S. Markus, and Anil M. Tuladhar. Network structure-function coupling and neurocognition in cerebral small vessel disease. NeuroImage: Clinical, 38, January 2023. Publisher: Elsevier Inc.

[27] Bo Hua, Xin Ding, Minghua Xiong, Fanyu Zhang, Yi Luo, Jurong Ding, and Zhongxiang Ding. Alterations of functional and structural connectivity in patients with brain metastases. PLoS ONE, 15(5), May 2020. Publisher: Public Library of Science.

[28] Angeliki Zarkali, Peter McColgan, Louise Ann Leyland, Andrew J. Lees, Geraint Rees, and Rimona S. Weil. Organisational and neuromodulatory underpinnings of structural-functional connectivity decoupling in patients with Parkinson’s disease. Communications Biology, 4(1), December 2021. Publisher: Nature Research.

[29] Guangyao Liu, Weihao Zheng, Hong Liu, Man Guo, Laiyang Ma, Wanjun Hu, Ming Ke, Yu Sun, Jing Zhang, and Zhe Zhang. Aberrant dynamic structure–function relationship of rich-club organization in treatment-naïve newly diagnosed juvenile myoclonic epilepsy. Human Brain Mapping, 43(12):3633–3645, August 2022. Publisher: John Wiley and Sons Inc.

[30] Ibai Diez, Paolo Bonifazi, Iñaki Escudero, Beatriz Mateos, Miguel A Muñoz, Sebastiano Stramaglia, and Jesus M Cortes. A novel brain partition highlights the modular skeleton shared by structure and function OPEN. Nature Publishing Group, 2015.

[31] Antonio Jimenez-Marin, Ibai Diez, Asier Erramuzpe, Sebastiano Stramaglia, Paolo Bonifazi, and Jesus M. Cortes. Open datasets and code for multi-scale relations on structure, function and neuro-genetics in the human brain. Scientific Data, 11(1), December 2024. Publisher: Nature Research.

[32] Richard F. Betzel, Maxwell A. Bertolero, Evan M. Gordon, Caterina Gratton, Nico U.F. Dosenbach, and Danielle S. Bassett. The community structure of functional brain networks exhibits scale-specific patterns of inter-and intra-subject variability. NeuroImage, 202, November 2019. Publisher: Academic Press Inc.

[33] Maria Grazia Puxeddu, Joshua Faskowitz, Olaf Sporns, Laura Astolfi, and Richard F. Betzel. Multi-modal and multi-subject modular organization of human brain networks. NeuroImage, 264, December 2022. Publisher: Academic Press Inc.

[34] Peter Fransson and Guillaume Marrelec. The precuneus/posterior cingulate cortex plays a pivotal role in the default mode network: Evidence from a partial correlation network analysis. NeuroImage, 42(3):1178–1184, September 2008.

[35] Shuai Huang, Jing Li, Liang Sun, Jieping Ye, Adam Fleisher, Teresa Wu, Kewei Chen, and Eric Reiman. Learning brain connectivity of Alzheimer’s disease by sparse inverse covariance estimation. NeuroImage, 50(3):935–949, April 2010.

[36] Mehdi Rahim, Bertrand Thirion, and Ga{\”e}l Varoquaux. Population shrinkage of covariance (PoSCE) for better individual brain functional-connectivity estimation. Medical Image Analysis, 54:138–148, May 2019. Publisher: Elsevier B.V.

[37] Kirsten L Peterson, Ruben Sanchez-Romero, Ravi D Mill, and Michael W Cole. Regularized partial correlation provides reliable functional connectivity estimates while correcting for widespread confounding. biorXiv, 2023.

[38] Raymond Salvador, John Suckling, Christian Schwarzbauer, and Ed Bullmore. Undirected graphs of frequency-dependent functional connectivity in whole brain networks. Philosophical Transactions of the Royal Society B: Biological Sciences, 360(1457):937–946, 2005. Publisher: Royal Society.

[39] Guillaume Marrelec, Alexandre Krainik, Hugues Duffau, Mélanie Pélégrini-Issac, Stéphane Lehéricy, Julien Doyon, and Habib Benali. Partial correlation for functional brain interactivity investigation in functional MRI. NeuroImage, 32(1):228–237, 2006. Publisher: Academic Press Inc.

[40] Srikanth Ryali, Tianwen Chen, Kaustubh Supekar, and Vinod Menon. Estimation of functional connectivity in fMRI data using stability selection-based sparse partial correlation with elastic net penalty. NeuroImage, 59(4):3852–3861, February 2012.

[41] G Varoquaux. Estimating brain functional connectivity and its variations from fMRI, 2019.

[42] Raphael Liégeois, Augusto Santos, Vincenzo Matta, Dimitri Van De Ville, and Ali H. Sayed. Revisiting correlation-based functional connectivity and its relationship with structural connectivity. Network Neuroscience, 4(4):1235–1251, November 2020. Publisher: MIT Press.

[43] Matthew R. Brier, Anish Mitra, John E. McCarthy, Beau M. Ances, and Abraham Z. Snyder. Partial covariance based functional connectivity computation using Ledoit-Wolf covariance regularization. NeuroImage, 121:29–38, November 2015. Publisher: Academic Press Inc.

[44] Usama Pervaiz, Diego Vidaurre, Mark W. Woolrich, and Stephen M. Smith. Optimising network modelling methods for fMRI. NeuroImage, 211, May 2020. Publisher: Academic Press Inc.

[45] Miguel Ibáñez-Berganza, Carlo Lucibello, Francesca Santucci, Tommaso Gili, and Andrea Gabrielli. Noise cleaning the precision matrix of short time series. Physical Review E, 108(2), August 2023. Publisher: American Physical Society.

[46] Jerome Friedman, Trevor Hastie, and Robert Tibshirani. Sparse inverse covariance estimation with the graphical lasso. Biostatistics, 9(3):432–441, July 2008.

[47] Olivier Ledoit and Michael Wolf. Honey, I Shrunk the Sample Covariance Matrix. The Journal of Portfolio Management, 30(4):110 – 119, 2004.

[48] Ga{\”e}l Varoquaux and R. Cameron Craddock. Learning and comparing functional connectomes across subjects. NeuroImage, 80:405–415, October 2013.

[49] Shuai Huang, Jing Li, Liang Sun, Jun Liu, Teresa Wu, Kewei Chen, Adam Fleisher, Eric Reiman, and Jieping Ye. Learning Brain Connectivity of Alzheimer’s Disease from Neuroimaging Data. Curran Associates, Inc., 2009.

[50] Amanda F. Mejia, Mary Beth Nebel, Anita D. Barber, Ann S. Choe, James J. Pekar, Brian S. Caffo, and Martin A. Lindquist. Improved estimation of subject-level functional connectivity using full and partial correlation with empirical Bayes shrinkage. NeuroImage, 172:478–491, May 2018. Publisher: Academic Press Inc.

[51] Stephen M. Smith, Karla L. Miller, Gholamreza Salimi-Khorshidi, Matthew Webster, Christian F. Beckmann, Thomas E. Nichols, Joseph D. Ramsey, and Mark W. Woolrich. Network modelling methods for FMRI. NeuroImage, 54(2):875–891, January 2011.

[52] Ga{\”e}l Varoquaux, Alexandre Gramfort, Jean Baptiste Poline, and Bertrand Thirion. Brain covariance selection: better individual functional connectivity models using population prior. Advances in neural information processing systems, 23, August 2010.

[53] Antonio Jimenez-Marin, Ibai Diez, Asier Erramuzpe, Sebastiano Stramaglia, Paolo Bonifazi, and Jesus M Cortes. Brain Hierarchical Atlas 2 (BHA2), July 2023.

[54] Yashar Zeighami, Trygve E. Bakken, Thomas Nickl-Jockschat, Zeru Peterson, Anil G. Jegga, Jeremy A. Miller, Jay Schulkin, Alan C. Evans, Ed S. Lein, and Michael Hawrylycz. A comparison of anatomic and cellular transcriptome structures across 40 human brain diseases. PLoS Biology, 21(4), April 2023. Publisher: Public Library of Science.

[55] Carlo Nicolini, Giulia Forcellini, Ludovico Minati, and Angelo Bifone. Scale-resolved analysis of brain functional connectivity networks with spectral entropy. NeuroImage, 211, May 2020. Publisher: Academic Press Inc.

[56] MEJ Newman. Modularity and community structure in networks. Technical report, 2006.

[57] Nguyen Xuan Vinh, Julien Epps, and James Bailey. Information Theoretic Measures for Clusterings Comparison: Variants, Properties, Normalization and Correction for Chance. Journal of Machine Learning Research, 11:2837– 2854, 2010.

[58] Aric A Hagberg, Daniel A Schult, and Pieter J Swart. Exploring Network Structure, Dynamics, and Function using NetworkX. In Ga\textbackslash”el Varoquaux, Travis Vaught, and Jarrod Millman, editors, Proceedings of the 7th Python in Science Conference, pages 11–15, Pasadena, CA USA, 2008.

[59] P. Van Mieghem, X. Ge, P. Schumm, S. Trajanovski, and H. Wang. Spectral graph analysis of modularity and assortativity. Physical Review E -Statistical, Nonlinear, and Soft Matter Physics, 82(5), November 2010.

[60] Fabian Pedregosa, Ga{\”e}l Varoquaux, Alexandre Gramfort, Vincent Michel, Bertrand Thirion, Olivier Grisel, Mathieu Blondel, Peter Prettenhofer, Ron Weiss, Vincent Dubourg, Jake Vanderplas, Alexandre Passos, David Cournapeau, Matthieu Brucher, Matthieu Perrot, and \’Edouard Duchesnay. Scikit-learn: Machine Learning in Python. Journal of Machine Learning Research, 12:2825–2830, 2011.

[61] Simone Romano, Nguyen Xuan Vinh, James Bailey, and Karin Verspoor. Adjusting for Chance Clustering Comparison Measures. Journal of Machine Learning Research, 17:1–32, 2016.

[62] Joël Bun, Jean-Philippe Bouchaud, and Marc Potters. Cleaning large correlation matrices: Tools from Random Matrix Theory. Physics Reports, 666:1–109, January 2017.

[63] Bernadette C. M. van Wijk, Cornelis J. Stam, and Andreas Daffertshofer. Comparing Brain Networks of Different Size and Connectivity Density Using Graph Theory. PLOS ONE, 5(10):e13701, 2010. Publisher: Public Library of Science.

[64] P. Read Montague, Raymond J. Dolan, Karl J. Friston, and Peter Dayan. Computational psychiatry. Trends in Cognitive Sciences, 16(1):72–80, January 2012.

[65] Klaas Enno Stephan and Christoph Mathys. Computational approaches to psychiatry. Current Opinion in Neurobiology, 25:85–92, April 2014.

[66] Karl J. Friston, Klaas Enno Stephan, Read Montague, and Raymond J. Dolan. Computational psychiatry: the brain as a phantastic organ. The Lancet. Psychiatry, 1(2):148–158, July 2014.

[67] Quentin J. M. Huys, Tiago V. Maia, and Michael J. Frank. Computational psychiatry as a bridge from neuroscience to clinical applications. Nature Neuroscience, 19(3):404–413, March 2016.

[68] Stefan Frässle, Yu Yao, Dario Schöbi, Eduardo A. Aponte, Jakob Heinzle, and Klaas E. Stephan. Generative models for clinical applications in computational psychiatry. Wiley Interdisciplinary Reviews. Cognitive Science, 9(3):e1460, May 2018.

[69] L. Rigoux and J. Daunizeau. Dynamic causal modelling of brain-behaviour relationships. NeuroImage, 117:202– 221, August 2015.

[70] Stefan Frässle, Zina M. Manjaly, Cao T. Do, Lars Kasper, Klaas P. Pruessmann, and Klaas E. Stephan. Whole-brain estimates of directed connectivity for human connectomics. NeuroImage, 225:117491, January 2021.

[71] Pauline Belujon and Anthony A Grace. Dopamine System Dysregulation in Major Depressive Disorders. International Journal of Neuropsychopharmacology, 20(12):1036–1046, June 2017.

[72] Edna Grünblatt, Tobias U. Hauser, and Susanne Walitza. Imaging genetics in obsessive-compulsive disorder: linking genetic variations to alterations in neuroimaging. Progress in Neurobiology, 121:114–124, October 2014.

[73] Tadafumi Kato. Current understanding of bipolar disorder: Toward integration of biological basis and treatment strategies. Psychiatry and Clinical Neurosciences, 73(9):526–540, September 2019.

[74] Sacha B. Nelson and Vera Valakh. Excitatory/Inhibitory Balance and Circuit Homeostasis in Autism Spectrum Disorders. Neuron, 87(4):684–698, August 2015.

[75] Javier Rasero, Antonio Jimenez-Marin, Ibai Diez, Roberto Toro, Mazahir T. Hasan, and Jesus M. Cortes. The Neurogenetics of Functional Connectivity Alterations in Autism: Insights From Subtyping in 657 Individuals. Biological Psychiatry, 94(10):804–813, November 2023.

[76] Britta E. Lindquist, Clare Timbie, Yuliya Voskobiynyk, and Jeanne T. Paz. Thalamocortical circuits in generalized epilepsy: Pathophysiologic mechanisms and therapeutic targets. Neurobiology of disease, 181:106094, June 2023.

[77] J Vuong and A. Devergnas. The Role of the Basal Ganglia in the Control of Seizure. Journal of neural transmission (Vienna, Austria : 1996), 125(3):531–545, March 2018.

[78] Michael A. DeTure and Dennis W. Dickson. The neuropathological diagnosis of Alzheimer’s disease. Molecular Neurodegeneration, 14(1):32, August 2019.

[79] Emilia M. Gatto, Natalia González Rojas, Gabriel Persi, José Luis Etcheverry, Martín Emiliano Cesarini, and Claudia Perandones. Huntington disease: Advances in the understanding of its mechanisms. Clinical Parkinsonism & Related Disorders, 3:100056, 2020.

[80] Anahit Babayan, Miray Erbey, Deniz Kumral, Janis D. Reinelt, Andrea M.F. Reiter, Josefin Röbbig, H. Lina Schaare, Marie Uhlig, Alfred Anwander, Pierre Louis Bazin, Annette Horstmann, Leonie Lampe, Vadim V. Nikulin, Hadas Okon-Singer, Sven Preusser, André Pampel Christiane S. Rohr, Julia Sacher, Angelika Thöne-Otto, Sabrina Trapp, Till Nierhaus, Denise Altmann, Katrin Arelin, Maria Blöchl, Edith Bongartz, Patric Breig, Elena Cesnaite, Sufang Chen, Roberto Cozatl, Saskia Czerwonatis, Gabriele Dambrauskaite, Maria Dreyer, Jessica Enders, Melina Engelhardt, Marie Michele Fischer, Norman Forschack, Johannes Golchert, Laura Golz, C. Alexandrina Guran, Susanna Hedrich, Nicole Hentschel, Daria I. Hoffmann, Julia M. Huntenburg, Rebecca Jost, Anna Kosatschek, Stella Kunzendorf, Hannah Lammers, Mark E. Lauckner, Keyvan Mahjoory, Ahmad S. Kanaan, Natacha Mendes, Ramona Menger, Enzo Morino, Karina Näthe, Jennifer Neubauer, Handan Noyan, Sabine Oligschläger, Patricia Panczyszyn-Trzewik, Dorothee Poehlchen, Nadine Putzke, Sabrina Roski, Marie Catherine Schaller, Anja Schieferbein, Benito Schlaak, Robert Schmidt, Krzysztof J. Gorgolewski, Hanna Maria Schmidt, Anne Schrimpf, Sylvia Stasch, Maria Voss, Annett Wiedemann, Daniel S. Margulies, Michael Gaebler, and Arno Villringer. Data descriptor: A mind-brain-body dataset of MRI, EEG, cognition, emotion, and peripheral physiology in young and old adults. Scientific Data, 6, 2019. Publisher: Nature Publishing Groups.

[81] Ross D Markello, Aurina Arnatkeviciute, Jean-Baptiste Poline, Ben D Fulcher, Alex Fornito, and Bratislav Misic. Standardizing workflows in imaging transcriptomics with the abagen toolbox. eLIfe, 2021.

[82] Michael J. Hawrylycz, Ed S. Lein, Angela L. Guillozet-Bongaarts, Elaine H. Shen, Lydia Ng, Jeremy A. Miller, Louie N. Van De Lagemaat, Kimberly A. Smith, Amanda Ebbert, Zackery L. Riley, Chris Abajian, Christian F. Beckmann, Amy Bernard, Darren Bertagnolli, Andrew F. Boe, Preston M. Cartagena, M. Mallar Chakravarty, Mike Chapin, Jimmy Chong, Rachel A. Dalley, Barry David Daly, Chinh Dang, Suvro Datta, Nick Dee, Tim A. Dolbeare, Vance Faber, David Feng, David R. Fowler, Jeff Goldy, Benjamin W. Gregor, Zeb Haradon, David R. Haynor, John G. Hohmann, Steve Horvath, Robert E. Howard, Andreas Jeromin, Jayson M. Jochim, Marty Kinnunen, Christopher Lau, Evan T. Lazarz, Changkyu Lee, Tracy A. Lemon, Ling Li, Yang Li, John A. Morris, Caroline C. Overly, Patrick D. Parker, Sheana E. Parry, Melissa Reding, Joshua J. Royall, Jay Schulkin, Pedro Adolfo Sequeira, Clifford R. Slaughterbeck, Simon C. Smith, Andy J. Sodt, Susan M. Sunkin, Beryl E. Swanson, Marquis P. Vawter, Derric Williams, Paul Wohnoutka, H. Ronald Zielke, Daniel H. Geschwind, Patrick R. Hof, Stephen M. Smith, Christof Koch, Seth G.N. Grant, and Allan R. Jones. An anatomically comprehensive atlas of the adult human brain transcriptome. Nature, 489(7416):391–399, September 2012.

[83] Olivier Ledoit and Michael Wolf. A well-conditioned estimator for large-dimensional covariance matrices. Journal of Multivariate Analysis, 88(2):365–411, 2004. Publisher: Academic Press Inc.

[84] Olivier Ledoit and Sandrine Péché. Eigenvectors of some large sample covariance matrix ensembles. Probability Theory and Related Fields, 151(1):233–264, October 2011.

[85] Joel Bun, Romain Allez, Jean Philippe Bouchaud, and Marc Potters. Rotational invariant estimator for general noisy matrices. IEEE Transactions on Information Theory, 62(12):7475–7490, December 2016. Publisher: Institute of Electrical and Electronics Engineers Inc.

[86] Joel Bun, Jean Philippe Bouchaud, and Marc Potters. My Beautiful Laundrette: Cleaning Correlation Matrices for Portfolio Optimization. May 2016.

[87] Pauli Virtanen, Ralf Gommers, Travis E. Oliphant, Matt Haberland, Tyler Reddy, David Cournapeau, Evgeni Burovski, Pearu Peterson, Warren Weckesser, Jonathan Bright, Stéfan J. van der Walt, Matthew Brett, Joshua Wilson, K. Jarrod Millman, Nikolay Mayorov, Andrew R. J. Nelson, Eric Jones, Robert Kern, Eric Larson, C. J. Carey, İlhan Polat, Yu Feng, Eric W. Moore, Jake VanderPlas, Denis Laxalde, Josef Perktold, Robert Cimrman, Ian Henriksen, E. A. Quintero, Charles R. Harris, Anne M. Archibald, Antônio H. Ribeiro, Fabian Pedregosa, and Paul van Mulbregt. SciPy 1.0: fundamental algorithms for scientific computing in Python. Nature Methods, 17(3):261–272, March 2020. Publisher: Nature Publishing Group.

[88] Irena Jovanović and Zoran Stanić. Spectral distances of graphs. Linear Algebra and Its Applications, 436(5):1425– 1435, March 2012.

[89] Carlo Lucibello. CarloLucibello/covariance-estimators, October 2023. original-date: 2021-09-30T14:49:37Z.

[90] Rossana Mastrandrea, Andrea Gabrielli, Fabrizio Piras, Gianfranco Spalletta, Guido Caldarelli, and Tommaso Gili. Organization and hierarchy of the human functional brain network lead to a chain-like core. Scientific Reports, 7(1):4888, July 2017. Publisher: Nature Publishing Group.

